# Dendritic Interaction of Timescales in Afterdepolarization Potentials and Nonmonotonic Spike-adding

**DOI:** 10.1101/2025.09.26.678799

**Authors:** Nils A. Koch, Yifan Zhao, Anmar Khadra

## Abstract

Depolarizations that occur after action potentials, known as afterdepolarization potentials or ADPs, are important for neuronal excitability and stimulus evoked transient bursting. Slow inward and fast outward currents underlie the generation of such ADPs with modulation of ADP amplitudes occurring as a result of neuronal morphology. However, the relative contribution and role of these slow inward and fast outward currents in ADP generation is poorly understood in the context of somatic and dendritic localization as well as with varied dendritic properties. Using a two compartmental Hodgkin-Huxley type model, the role of somatic and dendritic compartmentalization of ADP associated currents is investigated, revealing that dendritic (rather than somatic) slow inward and fast outward currents are the main contributors to ADP and spike-adding during both brief step current and AMPA current input. Additionally, dendritic size and passive properties of the dendrites were found to be key modulators of ADP amplitude. However, increasing magnitudes of NMDA current conductance resulted in nonmonotonic spike-adding in a manner dependent on dendritic Ca^2+^ influx and Ca^2+^ activated K^+^ currents, which was found to be the result of tight regulation of stimulus evoked transient bursting through positive feedback on action potential generation by dendritic Ca^2+^ and subsequent negative feedback through Ca^2+^ activated K^+^ currents. This novel mechanism of ADP and spike-adding regulation highlights the role of currents with slow timescales in ADPs, stimulus evoked bursting and neuronal excitability with implications for Ca^2+^ dependent synaptic plasticity and neuromodulation.

## 1 Introduction

In neurons across the central and peripheral nervous system, subthreshold depolarizations, termed afterdepolarization potentials (ADPs), can occur after action potentials (APs), also called transient burst, and thus modulate neuronal excitability and firing patterns (Gray and McCormick 1996; Amir et al. 2002; Chapin and Andrade 2000; Chen and Yaari 2008; Yue and Yaari 2006; D’Ascenzo et al. 2009; Connors et al. 1982; Greene et al. 1994; Staff et al. 2000; Li and Hatton 1997; Hilaire et al. 2005; Grampp 1966; Mayer 1985; Lemon and Turner 2000; Sheasby and Fohlmeister 1999; Stern and Armstrong 1996; Chu and Moenter 2006; Norekian 1999). ADPs enable neurons to store information on short timescales and may be important for working memory (Lisman and Idiart 1995; Klinshov and Nekorkin 2008; Fransén et al. 2002) as well as for temporal integration of inputs (Okamoto and Fukai 2009). Using single-compartment Hodgkin-Huxley models, it was shown that generating an ADP requires that the ionic currents underlying ADPs be slower than those driving action potentials (APs) and occur in such a way that the total post-AP inward current exceeds the total post-AP outward current (Nowacki et al. 2011). In particular, the timescale of this inward current must be slower than that of the outward current (Nowacki et al. 2011), for ADPs to form. Consequently, the magnitude of these fast outward and slow inward currents were found to be the primary drivers of ADPs including modulating ADP amplitude and altering the number of spikes within transient bursts (Nowacki et al. 2013).

ADPs can induce a temporal lowering of the firing threshold and as the ADP amplitude increases, this lowered neuronal firing threshold can be readily crossed resulting in an additional evoked AP, a phenomenon known as spike-adding (Klinshov and Nekorkin 2012). The transition from single evoked AP to the elicitation of a secondary AP was shown to be associated with saddle-node bifurcation of periodic orbits (Klinshov and Nekorkin 2012); however, spike-adding associated with ADP can also occur with a canard-like transition as a result of time-scale separation in the biophysical model distinct from periodic bursting but that does not involve bifurcations (Nowacki et al. 2012). Additionally, transient bursts that occur in response to brief inputs resulting from spike-adding require spike generation mechanism with timescales that promote bistability and a slow outward current within optimal ranges of magnitude and activation gating (Golomb et al. 2006). ADP-associated spike-adding and bursting is present in many neurons including in CA1 and CA3 pyramidal neurons (Gu et al. 2005; Xu and Clancy 2008; Nowacki et al. 2011), inferior olive neurons (Xu and Clancy 2008; Sultan et al. 2024), dorsal root ganglion neurons (White et al. 1989), Purkinje cells (Niday and Bean 2021), pyramidal neurons in the electrosensory lateral line lobe of weakly electric fish (Lemon and Turner 2000; Doiron et al. 2003; Metzen et al. 2025; Akhshi et al. 2025, 2023), lobster stretch receptor neurons (Grampp 1966), and gonadotropin-releasing Hormone neurons (Liu and Herbison 2008; Lehnert and Khadra 2019; Moran et al. 2016). Some neurons exhibit smaller ADPs which are less likely to trigger bursts, but lead to slow irregular firing (Amir et al. 2002). Moreover, vulnerable dopaminergic neurons in the substantia nigra pars compacta in a mouse model of Parkinson’s disease display larger ADPs and increased excitability than the resilient neuronal population (Beaver and Evans 2025), suggesting a role of ADPs in disorders of hyperexcitability.

Although, fast outward and slow inward currents underlie ADPs and spike-adding (Nowacki et al. 2013), multiple specific currents have been implicated in ADPs. Additionally, current flow from dendrites to soma is associated with increase in excitability and ADP formation (Lemon and Turner 2000; Doiron et al. 2003; Mainen and Sejnowski 1996), suggesting location-dependent effects of such currents. With regards to fast outward currents, K^+^ channels participate in ADP generation as evidenced by the association of decreased input resistance with ADPs, increased ADP amplitude resulting from K^+^ channel block with TEA and increased extracellular K^+^ concentrations leading to bursting (Yoshimura et al. 1987; Hilaire et al. 2005; Yue and Yaari 2004). For example, D-type K^+^ current block increases ADP amplitude and the number of APs evoked by a brief 3 ms stimulus in rat CA1 pyramidal neurons (Metz et al. 2007). However, local application of the D-type K^+^ channel blocker to the apical dendrites potentiating ADP amplitude suggesting that dendritic K^+^ currents are involved in regulating ADPs (Metz et al. 2007). Furthermore, electrotonic excision of dendrite, where the difference between dendritic and somatic potentials is eliminated, reduces somatic ADP amplitudes including during potentiation of ADPs with D-type K^+^ channel blocker further implicating the interaction of the soma and the dendrites in shaping ADP generation (Metz et al. 2007). ADP amplitude has also been found to be regulated by the inactivation of A-type K^+^ (Kv4) channels with larger ADPs corresponding to larger proportions of inactivated channels (Sultan et al. 2024). Dendritic and not somatic A-type K^+^ currents regulated ADPs as localized 4-AP application to CA1 pyramidal neuron dendrites but not to their somas increases ADPs and results in multiple evoked APs (Magee and Carruth 1999; Yue and Yaari 2006). In cases where 4-AP has no direct effect on ADPs, it has been suggested that the lack of effect is due to the majority of K^+^ channels affected by this blocker being inactivated at resting membrane potentials (Yue and Yaari 2004).

K^+^ currents with slower timescales also impact ADPs. Pharmacological block of the M-current results in increased ADPs and multiple evoked APs by brief step current inputs in CA1 pyramidal cells, with M-current potentiators having the opposite effect (Yue and Yaari 2004). In particular, axo-somatic M-current regulates the size and duration of ADPs in these neurons to prevent bursts from occurring, whereas dendritic M-current increases the threshold for initiating dendritic Ca^2+^ spikes and the associated transient somatic bursts (Yue and Yaari 2006). Both M-currents and Ca^2+^ activated K^+^ (K(Ca)) currents, which have similar slow timescales, are important for transient burst termination in neurons (Yue and Yaari 2006). Specifically, K(Ca) currents have been implicated in hyperpolarizing phases prior to ADPs (i.e. post-AP local minima prior to ADPs) (Poolos and Johnston 1999) and in hyperpolarization after ADPs (Higashi et al. 1993). In some cases, however, K(Ca) block has no direct effect on ADPs or bursting, but rather has a facilitatory effect on the number of APs evoked in bursts elicited by blocking the M-current (Yue and Yaari 2004) suggesting a complementary role of K(Ca) current in regulating ADP-associated bursting. This role is corroborated by the emergence of ADPs in dopamine neurons in rat substantia nigra with SK K^+^ channel block (Ping and Shepard 1999), increase in ADP amplitudes and induction of bursting with K(Ca) channel block in mouse Purkinje cells (Niday and Bean 2021), and the opposition of the ADP generating current by K(Ca) currents in gonadotropin-releasing hormone neurons Chu and Moenter (2006). Additionally, modeling has shown that the effect of varying Ca^2+^ conductance on stimulus driven bursting in some neurons is dependent on K(Ca) conductance (Golomb et al. 2006).

With regard to the slow outward current underlying ADPs, Ca^2+^ current dependent/mediated ADPs occur in many neurons (Yoshimura et al. 1987; Norekian 1999; Wong and Prince 1981; Chu and Moenter 2006). For instance, AP induced Ca^2+^ entry enhances ADPs in CA1 pyramidal cells during brief step current input, with N-type, P-/Q-type and T-/R-type Ca^2+^ current blockers decreasing ADP amplitude (Chen and Yaari 2008). Furthermore, intracellular application of Ca^2+^ chelator BAPTA suppresses ADPs and eliminates the effects of Ca^2+^ channel blockers on ADPs, suggesting that increases in intracellular Ca^2+^ concentration by Ca^2+^ channels contributes to ADP generation (Chen and Yaari 2008). Specifically, T-type Ca^2+^ currents have been shown to underlie ADPs and spike-adding in some neurons (Xu and Clancy 2008; White et al. 1989), likely because the T-type Ca^2+^ currents are the largest inward currents in these neurons and thus regulate the total inward current (Nowacki et al. 2011). Interestingly, calbindin negative dopaminergic neurons in the substantia nigra pars compacta have ADP-associated dendritic Ttype Ca^2+^ channel mediated Ca^2+^ increases not present in calbindin positive neurons (Evans et al. 2017), implicating T-type channel subcellular location as an important factor in their modulation of ADPs. Likewise, high voltage activated (HVA) Ca^2+^ channels have also been implicated in ADPs (Metz 2005; Wu et al. 2004) and in dendritic Ca^2+^ spike plateaus that are associated with synaptically driven bursts (Dudai et al. 2021). Such spike-adding was found to be the result of increases in dendritic *g*_*HV A*_ in a Hodgkin-Huxley model of layer 5 pyramidal neurons (Dudai et al. 2021). Additionally, somatic, and not dendritic, Ca^2+^ currents in hippocampal neurons drive ADPs and bursting (Jung et al. 2001; Traub et al. 1991) further implicating the subcellular location of Ca^2+^ currents in ADPs and spike-adding. Particularly, activation of dendritic Ca^2+^ currents by back-propagating APs produce slow inward currents that can generate ADPs and bursts of APs (Magee and Carruth 1999). However, dendritic Ca^2+^ current block has been also shown to attenuate stimulus-evoked bursting (Yue and Yaari 2006; Williams and Stuarty 1999), suggesting multiple roles of dendritic Ca^2+^ currents in the modulation of ADPs and spike-adding.

Although somatic or dendritic localization of ion channels can impact ADP generation, the morphological properties of neurons affect ADPs as well. For example, retinal ganglion cells show a negative correlation between dendritic size and firing responses (Sheasby and Fohlmeister 1999), hypothalamic magnocellular neurosecretory cells exhibit a relationship between ADP incidence and physiological decreases in dendritic length (Stern and Armstrong 1996, 1998; Teruyama and Armstrong 2002), and in hypothalamic gonadotropin-releasing Hormone neurons, smaller dendritic lengths are associated with increased ADP amplitude and repetitive firing (Roberts et al. 2008). In a population of two compartmental Hodgkin-Huxley models with different dendritic geometries but common distribution of ion channels, ADP size was found to decrease with reductions in the somato-dendritic coupling conductance and the proportion of dendritic area to somatic area (Mainen and Sejnowski 1996). In contrast, truncation of apical dendrites in rat CA1 pyramidal neurons does not alter ADP and the overall behavior to a brief (3 ms) step current input (Golomb et al. 2006), suggesting diversity in the role of dendrites in ADP generation. Further work using models consisting of a somatic and a dendritic compartment suggests that the proportion of dendritic area and coupling conductance dictate firing dynamics, with stimulus evoked bursting occurring at intermediate coupling conductance and increased dendritic area (Pinsky and Rinzel 1994). In multi-compartmental models that have more realistic dendritic arbor, dendritic length and topology (van Elburg and van Ooyen 2010; van Ooyen et al. 2002; Krichmar et al. 2002), as well as input location (Traub et al. 1991) affect stimulus evoked (van Elburg and van Ooyen 2010) and intrinsic (van Ooyen et al. 2002; Krichmar et al. 2002; Traub et al. 1991) bursting. Interestingly, the role of somato-dendritic coupling and dendritic area on single neuron firing extends to feed-forward networks of two compartmental models where these factors modulate the inputoutput characteristics of individual neurons to alter network activity and the transmission of signals through the network (Gao et al. 2022). Taken together, the dendritic size and interaction between dendrite and soma critically modulate ADPs and stimulus evoked bursting.

Although the contribution of different ionic channels (and their distribution), morphology and dendritic compartmentalization to ADPs, spike-adding and stimulus evoked bursting have been examined in isolation, a comprehensive analysis of how these factors collectively contribute and interact is, to the best of our knowledge, lacking. In this study, we therefore investigate the influence of specific ionic currents and dendritic compartmentalization on ADPs, spike-adding and stimulus evoked transient bursts of action potentials using models of cerebellar stellate cells that can be induced to burst intrinsically in a HVA Ca^2+^, A-type K^+^ and K(Ca) current dependent manner (Farjami et al. 2020a). Specifically, we find that dendritic morphology and dendritic HVA Ca^2+^ and K^+^ currents modulate ADPs and spike-adding and that the interaction of dendritic HVA Ca^2+^ and K(Ca) currents result in non-monotonic spike-adding in the presence of NMDA current inputs.

## 2 Methods

### 2.1 Cerebellar stellate cell models

The one compartment Hodgkin-Huxley type model of cerebellar stellate cells (CSC) described in Farjami et al. (2020a) was used and contains Na^+^ (*I*_*Na*_), delayed rectifier (Kdr) K^+^ (*I*_*Kd*_), leak (*I*_*L*_), A-type K^+^ (*I*_*A*_), T-type Ca^2+^ (*I*_*T*_), Ca^2+^ activated (K(Ca)) K^+^ (*I*_*K*(*Ca*)_), high voltage activated (HVA) Ca^2+^ (*I*_*HV A*_), and synaptic input (*I*_*syn*_) currents; it is given by

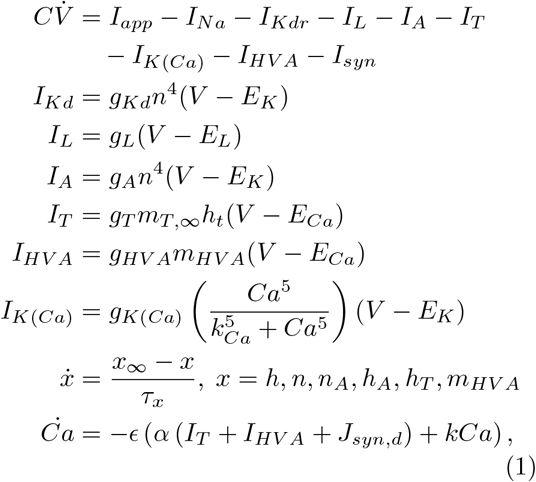

where *Ca* is cytosolic Ca^2+^ concentration, *J*_*syn*_ is the flux of Ca^2+^ due to the synaptic current in µM/ms, *C* = 1.50148 µF.cm^*−*2^ is the membrane capacitance and 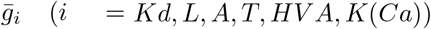 are the maximal conductances detailed in Table 1. Steady state gating of the currents is defined by Boltzmann functions of form 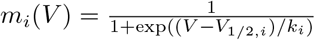 and are detailed along with gating time constants in Table 2. Additional model parameters are listed in Table 3.

**Table 1.** Parameter values of ionic currents for the CSC models. Conductances of the one compartment, along with the somatic and dendritic compartments of the two-compartment models are shown separately.

|  | One<br>Compartment<br>Model | Two<br>Compartment<br>Model |  |
| --- | --- | --- | --- |
| | $g$<br>($\mu\text{S}/\text{cm}^2$) | $g_{soma}$<br>($\mu\text{S}/\text{cm}^2$) | $g_{dendrites}$<br>($\mu\text{S}/\text{cm}^2$) |
| $I_{Na}$ | 3.4 | 3.4 | — |
| $I_{Kd}$ | 20.25 | 10.0112 | 10.2388 |
| $I_A$ | 12.2 | 6.8491 | 5.3509 |
| $I_T$ | 0.45045 | 0.2834 | 0.1670 |
| $I_{HVA}$ | 0.08 | 0.02047 | 0.05953 |
| $I_L$ | 0.07407 | 0.07407 | 0.07407 |
| $I_{K(Ca)}$ | 1.0 | 0.5505 | 0.4495 |

**Table 2.** Steady state activation and inactivation are defined as Boltzmann functions of form 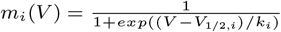 for each current. Time constants for activation and inactivation as well as reversal potentials are listed for each current. Values different for the dendritic compartment in the two compartmental model are listed in parentheses.

|  | Activation |  |  | Inactivation |  |  | E<br>(mV) |
| --- | --- | --- | --- | --- | --- | --- | --- |
| | $V_{1/2,x}$<br>(mV) | $k_x$ | $\tau_x$<br>(ms) | $V_{1/2,x}$<br>(mV) | $k_x$ | $\tau_x$<br>(ms) | |
| Na | -37 | 3 | — | -40 | -4 | $y0 + \frac{2Aw}{4\pi(V-V_c)^2+w^2}$ | 55 |
| Kd | -26 | 6 | $\frac{6}{1+e^{(V+23)/5}}$ | — | — | — | -80 |
| A | -27 | 13.2 | 5 | -82 | -6.5 | 10 | -80 |
| T | -54 | 3 | — | -74 | -3.75 | 15 | 22 |
| HVA | -25 | 8 | 4 | — | — | — | 22 |
| Leak | — | — | — | — | — | — | -38 (-60) |
| K(Ca) | $\frac{Ca^5}{k_{Ca}^5 + Ca^5}$ | | | — | — | — | -80 |

**Table 3.** Parameter Values for the CSC models.

|  |  |  |
| --- | --- | --- |
| $R_{axial}$ | 110 $\Omega \cdot \text{cm}$ | Rizza et al. (2021) |
| $r_{soma}$ | 10 $\mu\text{m}$ | — |
| $r_{cable}$ | 0.2 $\mu\text{m}$ | Abrahamsson et al. (2012) |
| $L$ | 150 $\mu\text{m}$ | — |
| $C_m$ | 1.50148 $\mu\text{F}/\text{cm}^2$ | Farjami et al. (2020b) |
| $g_{leak}$ | 0.07407 $\text{mS}/\text{cm}^2$ | Farjami et al. (2020b) |
| $R_m = 1/g_{leak}$ | 13.500742540839747 $\text{m}\Omega/\mu\text{F}$ | Farjami et al. (2020b) |
| $\Delta x$ | 15 $\mu\text{m}$ | — |
| $\kappa$ | 0.1163 | — |
| $\epsilon$ | 0.015 | Farjami et al. (2020b) |
| $\alpha$ | 0.018 $\mu\text{M cm}^2 \mu\text{A}^{-1} \text{ms}^{-1}$ | Farjami et al. (2020b) |
| $\alpha_s$ | 0.01538 $\mu\text{M cm}^2 \mu\text{A}^{-1} \text{ms}^{-1}$ | See Methods |
| $\alpha_d$ | 0.05182 $\mu\text{M cm}^2 \mu\text{A}^{-1} \text{ms}^{-1}$ | See Methods |
| $w$ | 46 $\text{mV}$ | Farjami et al. (2020b) |
| $V_c$ | -74 $\text{mV}$ | Farjami et al. (2020b) |
| $y_0$ | 0.1 $\text{ms}$ | Farjami et al. (2020b) |
| $A$ | 322 $\text{ms} \cdot \text{mV}$ | Farjami et al. (2020b) |
| $k$ | 0.1 $\text{ms}^{-1}$ | Farjami et al. (2020b) |
| $k_{Ca}$ | 0.45 $\mu\text{M}$ | Farjami et al. (2020b) |

Extension of this one compartment CSC model to a two compartmental model with somatic and dendritic compartments was done to assess the relative contributions of ion channels in the soma and dendrites of CSCs in shaping ADPs. The two compartmental model was developed using the formalism of Doiron et al. (2002), where the connection between the two compartments is modeled as resistive coupling with a coupling conductance *g*_*c*_ scaled by the ratio of somatic (*SA*_*soma*_) to total surface area of the somatic and dendritic (*SA*_*dend*_) compartments (*κ*) for the somatic compartment, i.e.,

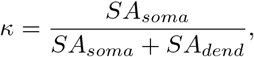

or scaled by the ratio 1 − *κ* for dendritic compartment. The two compartmental model is thus given by

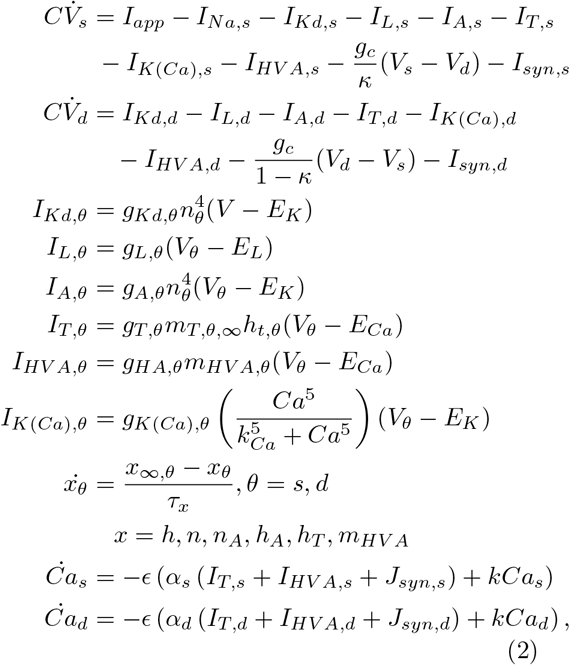

where 1/*g*_*c*_ is the axial resistance between the two compartments and *I*_*Na,θ*_, *I*_*Kd,θ*_, *I*_*L,θ*_, *I*_*A,θ*_, *I*_*T,θ*_, *I*_*K*(*Ca*),*θ*_, *I*_*HV a,θ*_, *I*_*syn,θ*_ are the Na^+^, Kdr, leak, A-type K^+^, T-type Ca^2+^, K(Ca), HVA Ca^2+^ and synaptic currents, respectively, in somatic (*θ* = *s*) and dendritic (*θ* = *d*) compartments. The currents from the one compartment model (Farjami et al. 2020a) were distributed to each compartment with the exception of the exclusively somatic presence of *I*_*Na*_. The maximum current conductances in each compartment were determined as the fraction of the conductance in the one compartment model attributable to each compartment as determined by the mean conductances presented in Rizza et al. (2021) (Table 1). The gating kinetics and reversal potentials (Table 2) as well as other model parameter values (Table 3) are unaltered from the one compartment model unless otherwise stated.

The dendritic compartment was chosen to be a cylinder with radius *r*_*d*_ and length *L*_*d*_ with one end connected to the soma. The somatic compartment is a sphere of radius *r*_*s*_ with the cap covered by the dendritic compartment missing from the surface area. The dendritic surface area is therefore 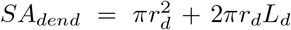 and the somatic surface area is 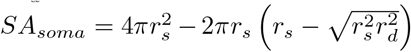, resulting in

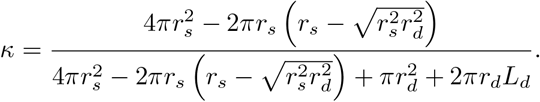

The conversion of Ca^2+^ current to the corresponding Ca^2+^ flux is 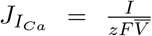 where *I* is the current that fluxes Ca^2+^ in µA, *z* is the valence of Ca^2+^ (*z* = 2), *F* is the Faraday con-stant, and 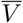 is the volume of the space into which the current fluxes. Given that model uses current density (µA/cm^2^), *I*_*Ca*_ must be multiplied by the surface area over which the channels exist resulting in the Ca^2+^ flux 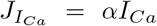 where 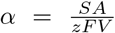.Thus the Ca^2+^ flux associated with somatic Ca^2+^ currents is 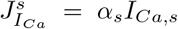 where 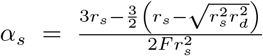 and the Ca^2+^ flux associated with dendritic Ca^2+^ currents is 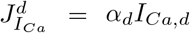 where 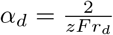.

### 2.2 Synaptic inputs

Synaptic input was generally simplified as a 3 ms long step current (Nowacki et al. 2013). More realistic AMPA current input approximated as a difference in exponentials, given by

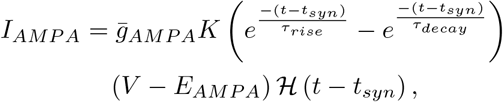

was used, where *g*_*AMP A*_ is the maximum conductance of excitatory synaptic input, *t*_*syn*_ is the time of onset of the synaptic input, ℋ(*t* − *t*_*syn*_) is the Heaviside step function and *K* is a normalization factor to ensure the difference of exponentials reaches a maximum of 1 at its peak (Carannante et al. 2022). Time constants appropriate for AMPA receptor synaptic input of *τ*_*rise*_ = 0.2 ms, and decay *τ*_*decay*_ = 2 ms (Kleppe and Robinson 1999) were used and the Ca^2+^ dynamics were altered to include the flux of Ca^2+^ associated with the AMPA current given the fractional AMPA Ca^2+^ current of 3% (*P*_*f,AMP A*_ = 0.03) (Burnashev et al. 1995) such that *J*_*syn,θ*_ = *α*_*θ*_*P*_*f,AMP A*_*I*_*AMP A*_ for synaptic input into compartment *θ*.

Calculation of *K* is dependent on the time of the maximal response *t*_*peak*_, given by

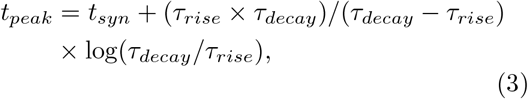

with

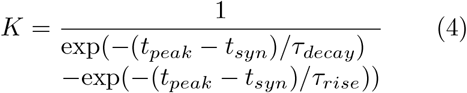

An NMDA current was also modeled as a difference of exponentials, but with the addition of the Mg^2+^ block (Wang 2002; Brunel and Wang 2001; Jahr and Stevens 1990), i.e.,

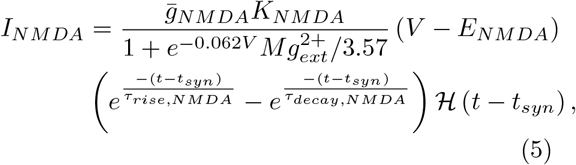

where 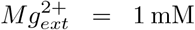 (Wang 2002; Brunel and Wang 2001), *τ*_*rise,NMDA*_ = 5 ms, and *τ*_*decay,NMDA*_ = 50 ms. The flux of Ca^2+^ associated with the NMDA current was accounted for by *J*_*syn,θ*_ = *α*_*theta*_*P*_*f,NMDA*_*I*_*NMDA*_, with a fractional Ca^2+^ current of 10% (*P*_*f,NMDA*_ = 0.1) (Burnashev et al. 1995; Watanabe et al. 2002; Burnashev 1996).

### 2.3 Quantification of afterdepolarization potentials

Peak detection was used to identify action potentials with a a minimum prominence of 25 mV. Detection of ADPs was limited to the time period after the last evoked AP by the synaptic input. ADPs were detected by the presence of at least two zero crossings of 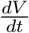) and at least three changes in inflection (i.e. zero crossings of 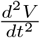) between the last evoked AP and the return to the holding membrane potential (Figure 1a-d) to ensure that only local maxima in membrane potential after AP were considered (Nowacki et al. 2013). The amplitude of each ADP was characterized in 2 ways: 1. The *amplitude* of the ADP peak above the holding potential. 2. The *relative amplitude* of the ADP peak above the local minima (defined by the zero crossing of 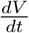) between the last AP and the ADP.

**Fig. 1.**
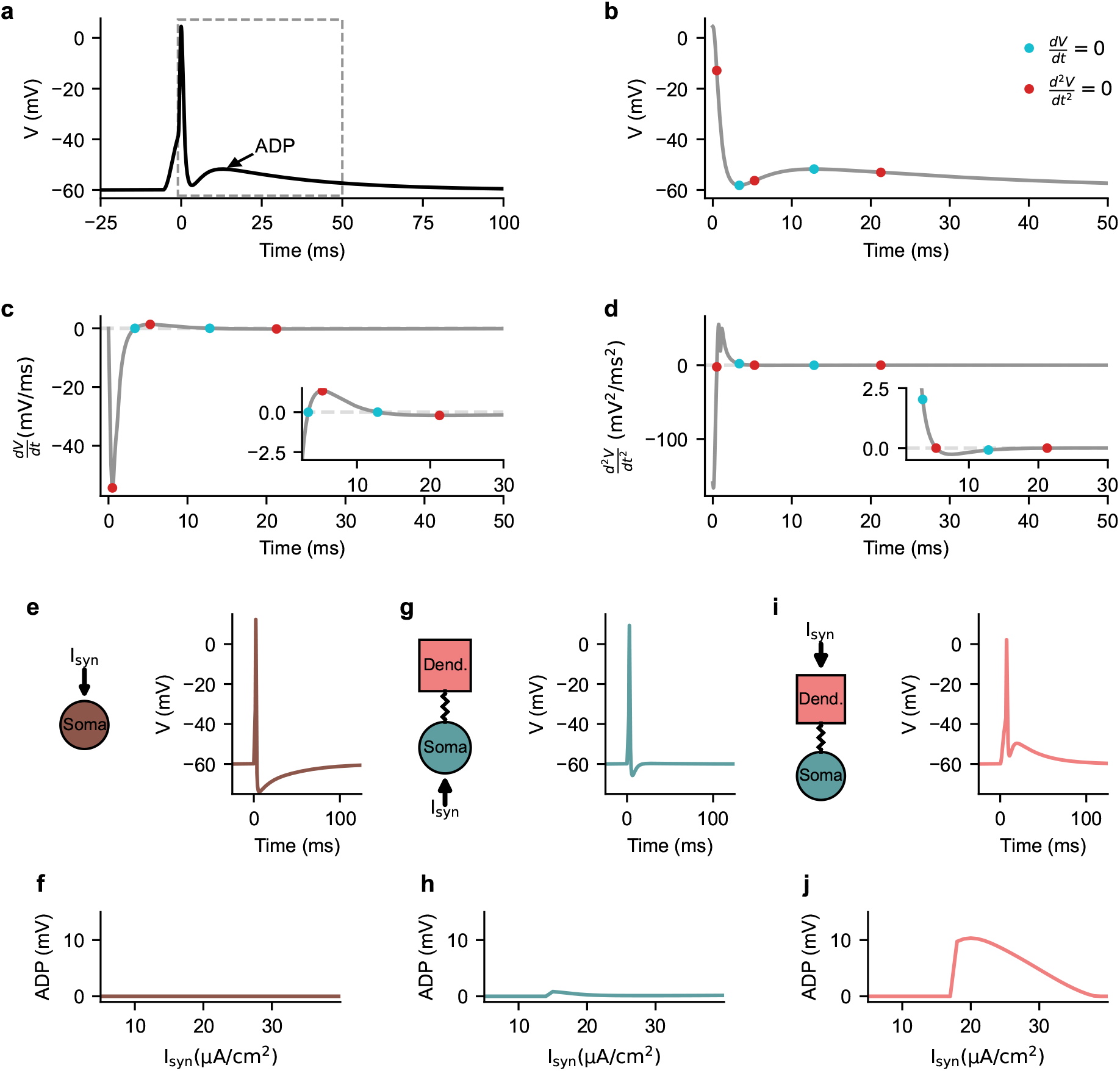
Afterdepolarizations and dendritic compartmentalization. (a) Example of somatic membrane potential response to a step current input displaying an afterdepolarization (ADP) following an action potential. (b) Zoomed-in region (gray dashed box in panel a) highlighting the ADP with first and second derivative null crossing marked in cyan and red, respectively. The first (c) and second (d) derivatives of somatic membrane potential corresponding to the time course in (b) with the null crossings shown (see insets). (e) Model schematic and the membrane potential response of the one compartment model to a 3 ms step current input. (f) The amplitude of the ADP elicited in response to different amplitudes of step current magnitudes (*I*_*syn*_) in the one compartment model. Schematic of the two compartmental model receiving a somatic (g) and dendritic (i) step current input (3 ms duration) and resulting somatic membrane potential response. The amplitude of the ADP elicited in response to different amplitudes of somatic (h) and dendritic (j) step current magnitudes (*I*_*syn*_) in the two compartmental model.

### 2.4 Sensitivity analysis

A variance-based sensitivity analysis using the Sobol method (Sobol 2001; Saltelli et al. 2008; Jansen 1999) modified from GlobalSensitivity.jl (Dixit and Rackauckas 2022) was performed. Quasi-Monte Carlo sampling was performed to sample the points in parameters space for the sensitivity analysis in a bounded manner with scrambling according to the method in Matoušek (1998) using QuasiMonteCarlo.jl. Specifically, conductances for slow inward (outward) currents were bounded between 0 and 5 (0.5 and 5) times their default values. Leak conductances were bounded between 0.5 and 1.5 times their default values, and the coupling conductance *g*_*c*_ was constrained between 0 and 2.5 times its default value. The ratio of somatic to total volume, *κ*, was constrained to values between 0 and 0.25. The magnitude of the step current was constrained to values between 0 and 50 µA/cm^2^.

### 2.5 Quantification of Ca^2+^ response and Ca^2+^ activated K^+^ current

As a measure of these signals, the cytosolic Ca^2+^ concentration and *I*_*K*(*Ca*)_ were integrated from the start of the synaptic input (*t*_*syn*_) to the time of the peak of the last evoked AP (*t*_*AP,last*_) using the trapezoidal rule on the simulated model trajectories and normalized by the number of evoked APs. i.e.,

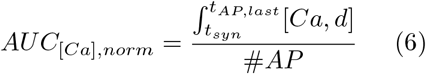

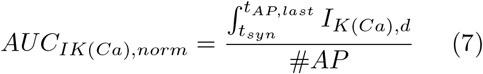

### 2.6 Simulation methods

Model simulations for all models were performed using the Julia programming language (Bezanson et al. 2017, Julia v1.10.0) with the Tsitouras 5/4 Runge-Kutta method implemented in DifferentialEquations.jl (Rackauckas and Nie 2017) starting at steady-state. A holding current was fit for each parameter set in each model to hold the somatic compartment at a holding potential of -60 mV. The fitting was performed by minimizing the squared difference of the steadystate somatic membrane potential with the desired holding potential using the NLopt (Johnson 2007) Subplex algorithm (Rowan 1990) with optimization using Optimization.jl (Dixit and Rackauckas 2023). To ensure global convergence, a sequential fitting approach is used, in which particle swarm optimization (Mogensen and Riseth 2018) was used whenever the output somatic membrane potential deviated by more than 0.1 mV from the desired holding potential, thereby ensuring that the steady-state holding potential of -60 mV was achieved across parameter sets. The simulation and analysis code is freely available online at https://github.com/nkoch1/ADP Koch et al.git.

## 3 Results

Afterdepolarization potentials (ADPs) were detected by the presence of at least two null crossings of the first voltage derivative and three null crossings of the second voltage derivative following the last action potential (AP) evoked (Figure 1a-d; see Methods). While a step current input into the one compartmental model evokes a single AP (Figure 1e), it does not evoke any ADPs regardless of the magnitude of the step current input (Figure 1f). In the two compartmental model, on the other hand, somatic step current input (Figure 1g) can evoke ADPs of small amplitudes (Figure 1h), whereas step current input into the dendritic compartment (Figure 1i) produces larger ADPs at sufficiently large input magnitudes (Figure 1j). Given that dendritic step current inputs are able to evoke larger ADPs across a greater range of input magnitudes, dendritic input in the two compartmental model will the main focus going forward.

### 3.1 Sensitivity analysis and subcellular contributions to ADPs

The contributions of model components to ADPs evoked with dendritic step current input (Figure 2a) on ADP amplitude, relative ADP amplitude and number of evoked APs (Figure 2b) was examined. Specifically, model parameters such as those relating to morphology (coupling conductance *g*_*c*_ and proportion of somatic surface area *κ*; Figure 2c-d), step current amplitude (Figure 2e), somatic current conductances (Figure 2f-j) and dendritic conductances (Figure 2k-o) were each investigated one at a time. In this context, the coupling conductance between somatic and dendritic compartments (Figure 2c), the relative somatic area (*κ*; Figure 2d), step current input magnitude (Figure 2e), somatic and dendritic A-type K^+^ conductance (Figure 2g and l, respectively) and dendritic HVA Ca^2+^ conductance (Figure 2n) were found to have the greatest impact on ADPs, with the dendritic HVA Ca^2+^ conductance causing the largest change to the number of evoked APs (Figure 2n).

**Fig. 2.**
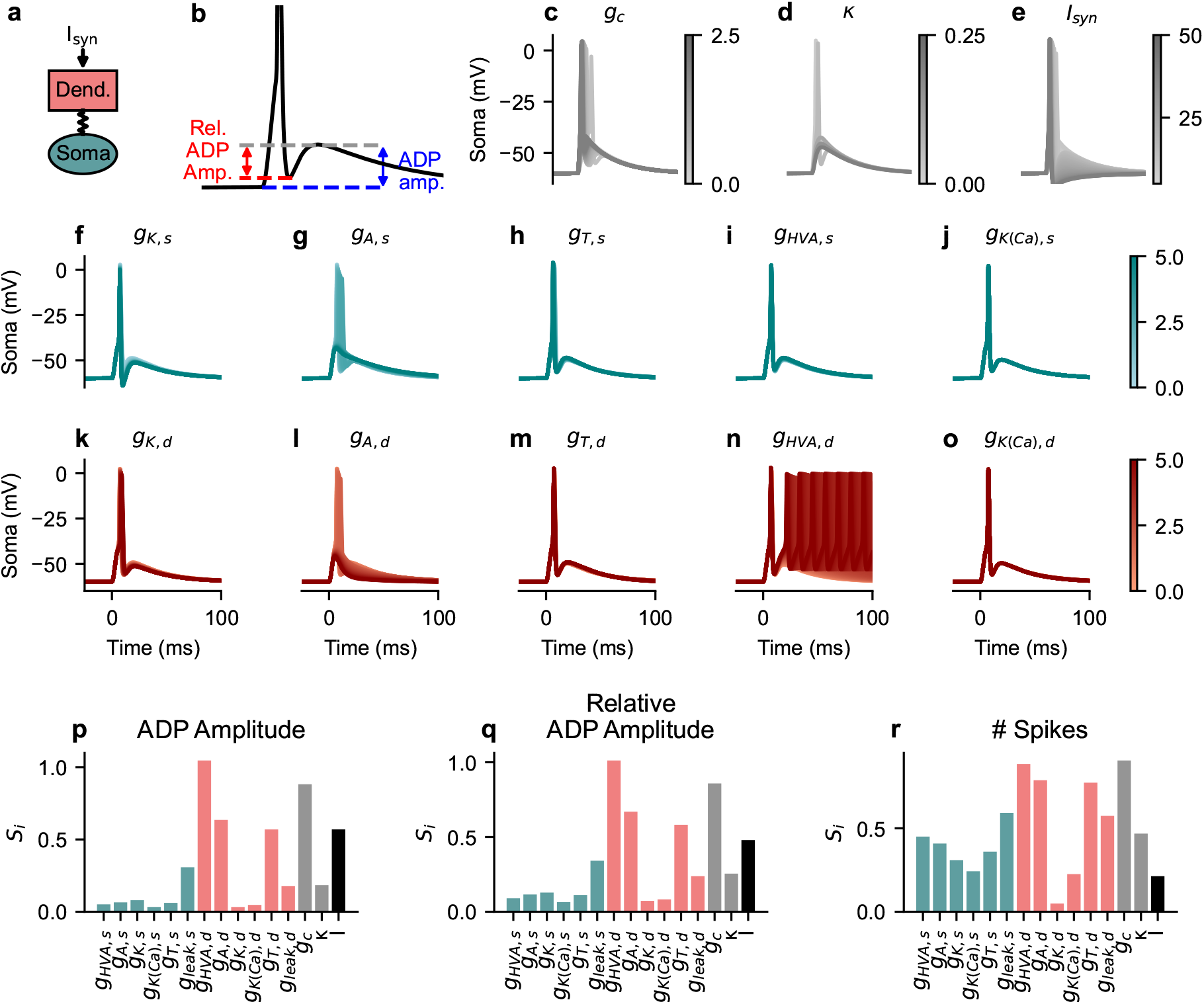
Sensitivity analysis of two compartmental model highlighting the role of dendritic currents. Using a model with somatic and dendritic compartments and dendritic step current input (a) ADPs can be evoked, and (b) quantified by their amplitude (blue) and relative amplitude from the preceding local minima (red). Effects of altering both the coupling conductance (*g*_*c*_; c),and the proportion of dendritic surface area (*κ*; d), as well as the magnitude of the input step current (e). The primary effect of scaling Kdr (*g*_*K,s*_; f), A-type K^+^ (*g*_*A,s*_; g), T-type Ca^2+^ (*g*_*T,s*_; h), HVA Ca^2+^ (*g*_*HV A,s*_; i) and K(Ca) (*g*_*K*(*Ca*),*s*_; j) conductances suggest negligible effect on ADPs. Scaling of dendritic Kdr (*g*_*K,d*_; k), A-type K^+^ (*g*_*A,d*_; l), T-type Ca^2+^ (*g*_*T,d*_; m), HVA Ca^2+^ (*g*_*HV A,d*_; n) and K(Ca) (*g*_*K*(*Ca*),*d*_; o) suggests greater involvement of dendritic currents. Variance-based sensitivity analysis formalized as Sobol indices for ADP amplitude (p), relative ADP amplitude (q) and the number of spikes evoked (r).

These qualitative observations across one parameter change at a time were then further examined by performing a variance-based sensitivity analysis using the Sobol method to quantify the effects of these model parameters on ADPs and spike-adding across parameter interactions.

The Sobol indices resulting from this method reflect the relative contribution of each model parameter on ADP amplitude (Figure 2p) and relative ADP amplitude (Figure 2q), indicating that dendritic Ca^2+^ (HVA and T-type) currents, A-type K^+^ current, coupling conductance between the two compartments and step current input magnitude are key modulators of ADP and relative ADP amplitude. While, these currents play an important role in determining the number of evoked APs, the leak conductances in both soma and dendritic compartments as well as somatic currents additionally contribute to the number of evoked APs (Figure 2r).

Interestingly, when applying somatic step current input, both dendritic and somatic currents, particularly dendritic HVA Ca^2+^, somatic and dendritic A-type K^+^ and leak currents were found to dictate ADPs and spike-adding (Figure S1); this is due to the propagation of the response signal from the soma to the dendrites rather than from the dendrites to the soma.

### 3.2 Effect of dendrites on ADPs

Previous work (Roberts et al. 2008; Sheasby and Fohlmeister 1999; Stern and Armstrong 1996, 1998; Teruyama and Armstrong 2002; Mainen and Sejnowski 1996) as well as the sensitivity analysis performed here (Figure 2) suggest that dendritic size can modulate ADP amplitudes. To this end, we investigated the role of dendritic size in the two compartmental model through the ratio of dendritic to total surface area of dendrites and soma, i.e., 1 − *κ*. The amplitude of ADPs, ADP amplitudes relative to the preceding local minima, and the number of APs evoked as 1 − *κ* increases was examined in combination with the magnitude of the step current input (Figure 3a), AMPA conductance (*g*_*AMP A*_; Figure 3b) and NMDA conductance (*g*_*NMDA*_; Figure 3c). Our results revealed that as the dendrites increase in size, ADPs emerge with modest spike-adding during both step current and AMPA current input (Figure 3a-b). However, during NMDA current input (in the presence of AMPA with *g*_*AMP A*_ = 10 µS/cm), spike-adding is more prevalent with greater number of APs added as 1 *− κ* increases (Figure 3c). Within a regime of 1 − *κ* and *g*_*NMDA*_ that elicits the same number of APs to the synaptic input, both ADP amplitude and the relative ADP amplitude increase as 1 − *κ* increases. Such a regime is flanked by the one eliciting one fewer AP and another eliciting one additional AP (Figure 3c), with ADP amplitude increasing toward the regime with an additional AP, highlighting the dependence of spike-adding on ADP amplitude.

**Fig. 3.**
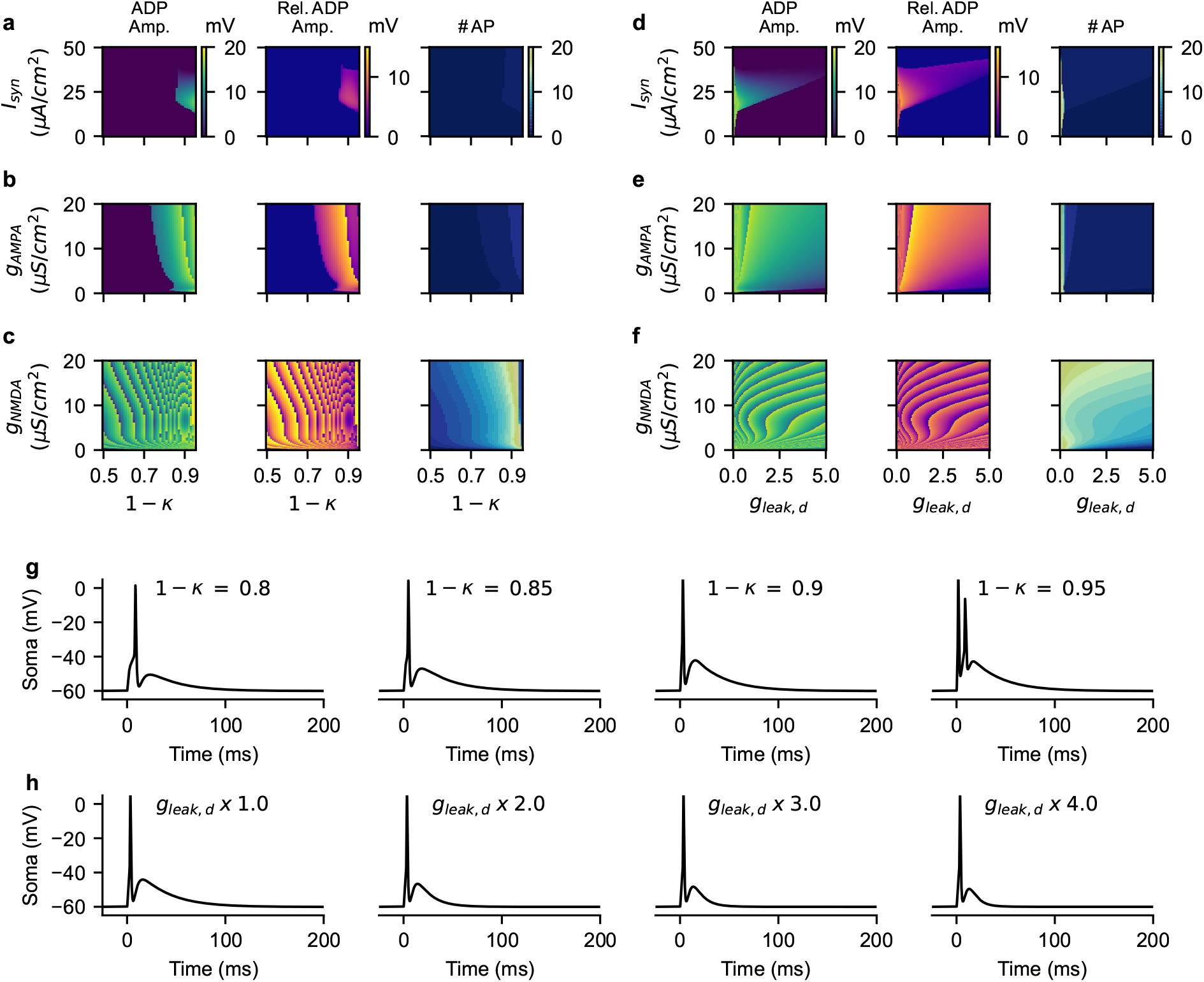
Effects of coupling and dendritic leak current on afterdepolarization. The effect of changing dendritic length, i.e., changing the ratio of dendritic to total surface area (1 *− κ*) of the two compartment model, on ADPs, evaluated in combination with varying step current input magnitude (*I*_*syn*_; a), AMPA current conductance (*g*_*AMP A*_; b) and NMDA current conductance (*g*_*NMDA*_; c) on ADP amplitude, relative ADP amplitude and number of spikes elicited. The effect of altering the dendritic leak conductance *g*_*leak,d*_ (in fold change from default value) on ADPs, evaluated in combination with varying step current input magnitude (d), AMPA current conductance (e) and NMDA current conductance (f). NMDA synaptic input occurred in the presence of an AMPA current with fixed conductance (*g*_*AMP A*_ = 10 µS/cm^2^). Example somatic voltage traces for increasing dendritic length (i.e. increasing 1 − *κ*; g) and for increasing dendritic leak current (h) at *g*_*AMP A*_ = 10 µS/cm^2^.

As the dendritic leak currents have been suggested to modulate ADP amplitudes (Roberts et al. 2008), the effects of altering the dendritic leak conductance (*g*_*leak,d*_ from 0 to 5 fold its default value on the amplitude of ADP, relative ADP amplitude and the number of APs evoked in concert with changes in the magnitude of the step current input (Figure 3d), AMPA conductance (*g*_*AMP A*_; Figure 3e) and NMDA conductance (*g*_*NMDA*_; Figure 3f) was investigated. For step current and AMPA dendritic inputs, we found that reduction of dendritic leak currents results in decreased ADP amplitudes and relative amplitude, with spike-adding occurring only when *g*_*leak,d*_ is close to zero (Figure 3d and e, respectively). In the presence of NMDA current, reduction of *g*_*leak,d*_ results in spike-adding with corresponding increases in both ADP and relative ADP amplitudes (Figure 3d), once again underscoring ADP involvement in spike-adding. This effect is illustrated in an example time-series simulations showing how increasing 1 − *κ* leads to progressive ADP growth and ultimately to spike-adding (Figure 3g). Conversely, simulations with increasing dendritic leak conductance (*g*_*leak,d*_) causes a reduction in ADP amplitude and relative amplitude (Figure 3h).

However, dendritic arbors are more complex than the single dendritic compartment considered in the model. This raises the question of whether approximating the entire arbor as a single compartment is reasonable in this context. To address this, we assessed ADP and spike adding dynamics using a ball-and-stick model in which the dendritic compartment is replaced with a dendritic cable (see Supplemental Material). The effect of dendritic leak (*g*_*leak,d*_) and dendritic size (1 − *κ*) in the two compartmental model (Figure S2a-c) is qualitatively similar to the effect of dendritic leak (*g*_*leak,d*_) and dendritic size in the ball-and-stick model (Figure S2d-f). Adding more dendritic branches to the model did not alter the effect of the dendritic leak *g*_*leak,d*_ compared to the original ball-and-stick model, nor did increasing the length of this additional dendritic cable alter ADP amplitude (Figure S2g-h). Thus, the approximation of the dendrites as a single compartment is a reasonable assumption and the two compartmental and ball-and-stick models both suggest that dendritic size as well as the passive properties of dendrites modulate ADP occurrence and amplitude and as such spike-adding in stimulus evoked neuronal responses.

### 3.3 Somatic and dendritic current modulation of ADPs

It has been previously shown that ADPs can be modulated by fast outward and slow inward currents (Nowacki et al. 2013). We therefore examined the influence of such currents in the somatic and dendritic compartment on ADPs and spike-adding in response to step current inputs; to do so, we focused on the effects of four key currents: the slow inward high voltage activated (HVA) Ca^2+^ current, the fast outward delayed rectifier (Kdr) current, the A-type K^+^ current and the Ca^2+^ activated K^+^ (K(Ca)) current. This was done initially by investigating the combined effects of scaling the somatic HVA conductance (*g*_*HV A,s*_) with the somatic A-type (Figure 4a; *g*_*A,s*_), with Kdr (Figure 4b; *g*_*Kd,s*_) and with K(Ca) (Figure 4c; *g*_*K*(*Ca*),*s*_) conductances, revealing that they have negligible effects on ADP amplitude and spike-adding. A-type and Kdr currents modulate the relative ADP amplitude from the preceding local minima (Figure 4b-c middle), but not the ADP amplitude itself (Figure 4b-c left), likely due to that fact that these currents shape the repolarization phase of the last AP and thus the local minima preceding the ADP rather than directly altering ADP amplitude. In contrast, scaling dendritic *g*_*HV A,d*_ with dendritic *g*_*A,d*_ (Figure 4d), with *g*_*Kd,d*_ (Figure 4e) and with *g*_*K*(*Ca*),*d*_ (Figure 4f) all produce qualitatively similar modulation of ADP amplitude and spike-adding. Specifically, ADP amplitudes and relative amplitude increase as *g*_*HV A,d*_ increases until a spike-adding regime is entered during which ADP amplitudes increase until another AP is added. Interestingly, at high dendritic *g*_*HV A,d*_ the dendritic A-type K^+^ current at large magnitudes can curtail spike-adding (upper right of Figure 4d right) whereas increases in dendritic *g*_*Kd,d*_ (Figure 4e) and *g*_*K*(*Ca*),*d*_ (Figure 4f) can reduce the number of evoked APs but not prevent spike-adding. Furthermore, increasing *g*_*A,d*_ increases the number of evoked APs until spike-adding stops (Figure 4d), whereas increasing *g*_*K*(*Ca*),*d*_ decreases the number of evoked APs steadily (Figure 4f). This likely reflects that the A-type K^+^ current has a faster timescale than the K(Ca) current. The increase in ADP amplitude and spike-adding with increasing *g*_*HV A,d*_ is demonstrated by the example traces in Figure 4g. When the T-type Ca^2+^ current is considered instead of the HVA Ca^2+^ current, no spike-adding is seen and modulation of ADP amplitudes is modest (Figure S3), which may reflect the faster timescale of T-type Ca^2+^ current compared to the HVA Ca^2+^ current. In general, dendritic and not somatic slow-inward and fast-outward currents shape ADP amplitudes and spike-adding to brief step current inputs.

**Fig. 4.**
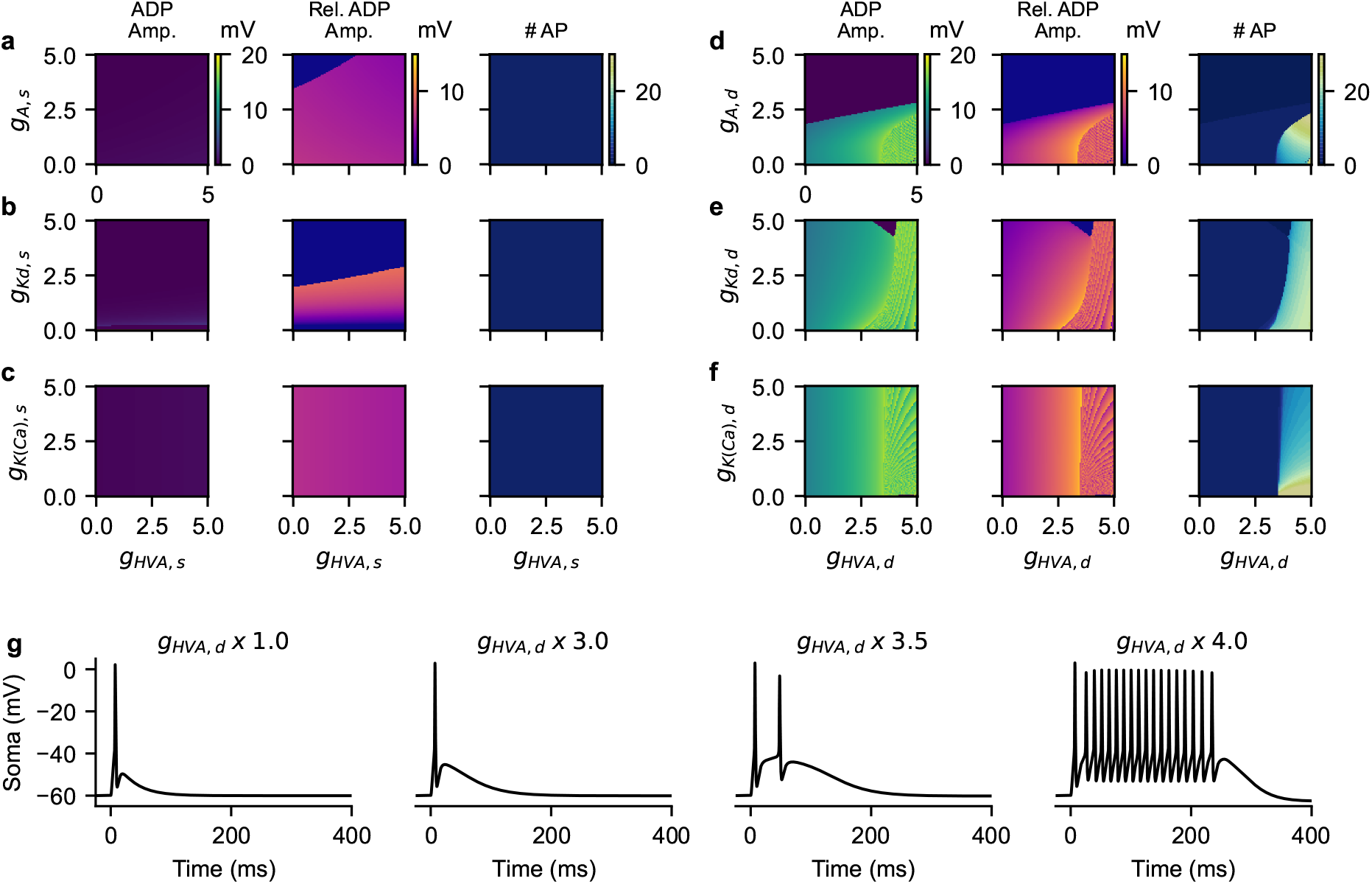
Spike-adding associated with the Dendritic Ca^2+^ and K^+^ currents evoked by step current input. The effect of altering somatic A-type K^+^ (a; *g*_*A,s*_), Kdr (b; *g*_*Kd,s*_) and K(Ca) (c; *g*_*K*(*Ca*),*s*_) current conductances in combination with varying somatic HVA Ca^2+^ (*g*_*HV A,s*_) current conductance during 3 ms step current input to the dendrite (20 µA/cm^2^) on ADP amplitude, relative ADP amplitude and the number of spikes evoked. The effect of altering dendritic A-type K^+^ (d; *g*_*A,d*_), Kdr (e; *g*_*Kd,d*_) and K(Ca) (f; *g*_*K*(*Ca*),*d*_) in combination with varying dendritic HVA Ca^2+^ (*g*_*HV A,d*_) during step current input to the dendrite (identical in magnitude to that in a, b and c) on ADP amplitude, relative ADP amplitude and the number of spikes evoked. The somatic voltage responses (g) displaying spike-adding seen with increasing dendritic HVA Ca^2+^ conductance in d-f in response to the same step current input. All conductance values are reported in fold change from default value.

As with the step current input, scaling somatic HVA conductance (*g*_*HV A,s*_) with somatic A-type (Figure 5a; *g*_*A,s*_), with Kdr (Figure 5b; *g*_*Kd,s*_) and with K(Ca) (Figure 5c; *g*_*K*(*Ca*),*s*_) conductances during AMPA current input had minimal effects on ADP amplitude and spike-adding. Additionally, *g*_*A,s*_ and *g*_*Kd,s*_ modulated the relative ADP amplitude from the preceding local minima (Figure 5b-c middle), but not ADP amplitude (Figure 5b-c left) in line with the effects observed with step current inputs, highlighting the role of these currents in shaping the repolarization of the last AP rather than ADP amplitude. Scaling the dendritic *g*_*HV A,d*_ with dendritic *g*_*A,d*_ (Figure 5d), with *g*_*Kd,d*_ (Figure 5e) and with *g*_*K*(*Ca*),*s*_ (Figure 5f) during AMPA current input produce qualitatively similar effects on ADP and spike-adding to that seen with step current input (Figure 4d-f), namely, that spike-adding generally occurs at larger *g*_*HV A,d*_ conductances. The spike-adding induced by increasing *g*_*HV A,d*_ during AMPA current input is demonstrated by the example traces in Figure 5g and is similar in nature to that seen during step current input (Figure 4g).

**Fig. 5.**
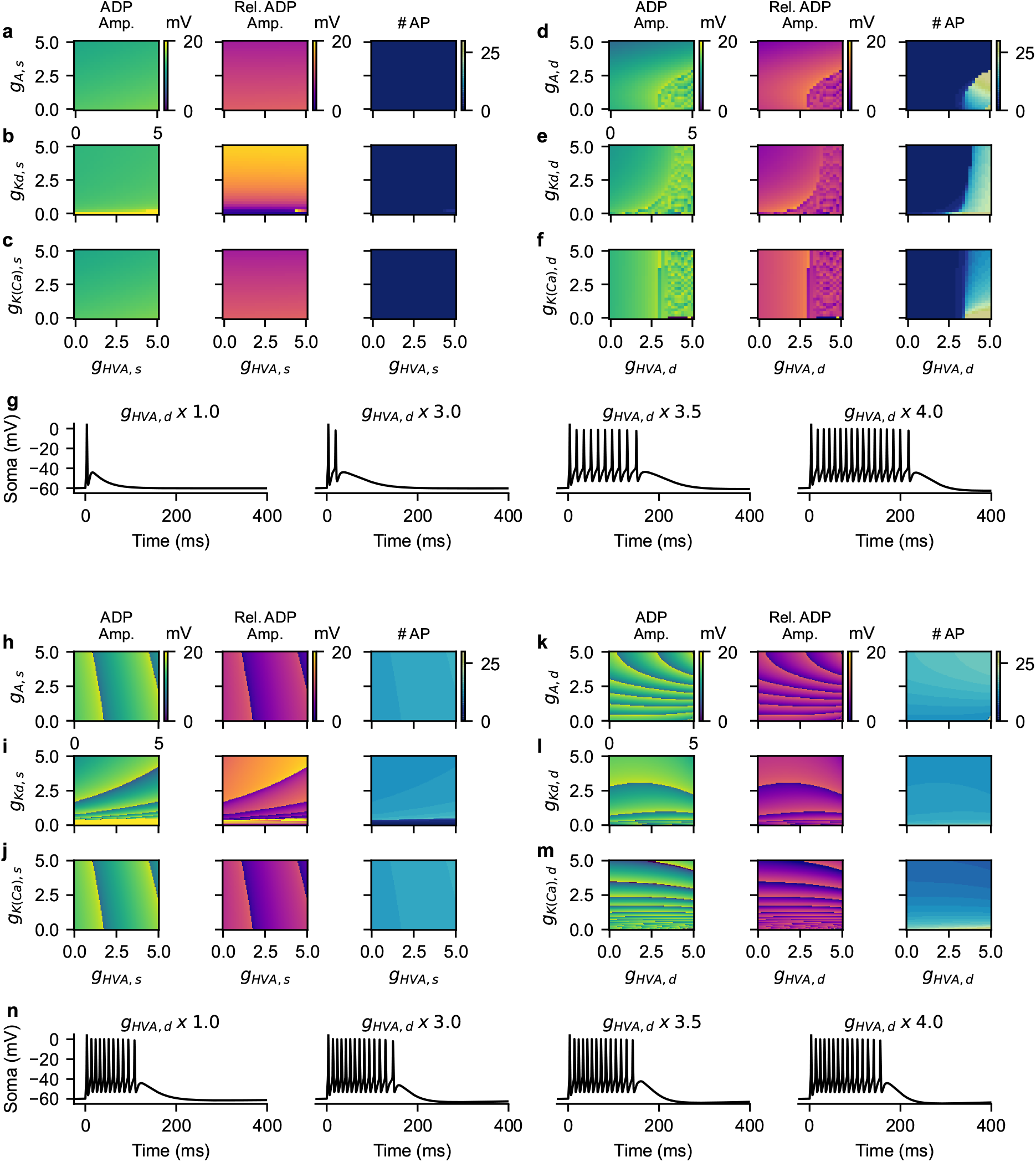
Dendritic Ca^2+^ and K^+^ currents modulate spike-adding mediated by AMPA and NMDA current inputs. The effect of altering somatic A-type K^+^ (a; *g*_*A,s*_), Kdr (b; *g*_*Kd,s*_) and K(Ca) (c; *g*_*K*(*Ca*),*s*_) current conductances in combination with varying somatic HVA Ca^2+^ (*g*_*HV A,s*_) current conductance on ADP amplitude, relative ADP amplitude and the number of spikes evoked in the presence of AMPA current input (*g*_*AMP A*_ = 10 µS/cm^2^). The effect of altering dendritic A-type K^+^ (d;*g*_*A,d*_), Kdr (e; *g*_*Kd,d*_) and K(Ca) (f; *g*_*K*(*Ca*),*d*_) current in concert with varying dendritic HVA Ca^2+^ (*g*_*HV A,d*_) current on ADP amplitude, relative ADP amplitude and the number of spikes evoked during this AMPA current input. Somatic voltage responses (g) showcasing spike-adding with increasing dendritic HVA Ca^2+^ conductance (*g*_*HV A,d*_). Effects of somatic A-type K^+^ (h; *g*_*A,s*_), Kdr (i; *g*_*Kd,s*_) and K(Ca) (j; *g*_*K*(*Ca*),*s*_) currents in concert with varying somatic HVA Ca^2+^ (*g*_*HV A,s*_) current, as well as the effects of dendritic A-type K^+^ (k; *g*_*A,d*_), Kdr (l; *g*_*Kd,d*_) and K(Ca) (m; *g*_*K*(*Ca*),*d*_) while altering dendritic HVA Ca^2+^ (*g*_*HV A,d*_) on ADP amplitude, relative ADP amplitude and the number of spikes evoked after the addition of an NMDA current to the AMPA current (*g*_*NMDA*_ = 2 µS/cm^2^). Somatic voltage responses showcasing how (n) an AMPA and NMDA current evokes numerous action potentials across increasing values of dendritic HVA Ca^2+^ conductance (*g*_*HV A,d*_). All conductance values are reported in fold change from the default value.

Unlike during step current and AMPA current inputs, scaling of somatic HVA conductance (*g*_*HV A,s*_) with somatic A-type (Figure 5h; *g*_*A,s*_), with Kdr (Figure 5i; *g*_*Kd,s*_) and with K(Ca) (Figure 5j; *g*_*K*(*Ca*),*s*_) conductances during NMDA current input modulates both ADP amplitudes and the number of evoked APs. In particular, increases in *g*_*HV A,s*_ dominate ADP amplitude and spike-adding modulation, whereas *g*_*A,s*_ (Figure 5h) had lesser impact as evidenced by the vertical nature of the boundaries between regions with different numbers of evoked APs. However, the somatic Kdr conductance (*g*_*Kd,s*_) contributed more strongly to ADP amplitude and spike-adding than the other somatic K^+^ currents as reflected by the less vertical increases in ADP amplitude and spike-adding (Figure 5i). The effect of *g*_*K*(*Ca*),*s*_ (Figure 5j) is less than that of *g*_*HV A,s*_ with more vertical boundaries between regions with different number of evoked APs in a manner similar to *g*_*A,s*_ (Figure 5h). Scaling dendritic HVA conductance *g*_*HV A,d*_ with dendritic A-type *g*_*A,d*_ (Figure 5k), with Kdr *g*_*Kd,d*_ (Figure 5l) and with K(Ca) *g*_*K*(*Ca*),*s*_ (Figure 5m) conductances further modulates ADPs and spike-adding during NMDA current input, with more impact on ADP amplitude, relative amplitude and number of evoked APs along the K^+^ current conductances axes than seen with somatic scaling. This is in line with the increased contribution of dendritic K^+^ currents on ADPs and spike-adding with step current (Figure 4 or only AMPA current inputs (Figure 5a-f). The modulation in the number of evoked APs with increasing *g*_*HV A,d*_ during AMPA and NMDA current inputs is demonstrated by the traces in Figure 5n. Additionally, modulation of T-type and not HVA Ca^2+^ current, does not result in spike-adding in the presence of AMPA current (Figure S3a-g) similar to during step current input; however, dendritic but not somatic T-type Ca^2+^ current modulation affects the number of evoked APs and ADP amplitudes (Figure S4h-n) during NMDA current input, suggesting that the T-type Ca^2+^ current on its own is insufficient to evoke spike-adding in the absence of a slow inward process such as an NMDA current. This may be a result of the timescale of the T-type Ca^2+^ current not being sufficiently slow. Taken together, these results indicate that dendritic Ca^2+^ and K^+^ current modulation of ADP amplitudes and spike-adding behavior is larger than the corresponding somatic currents during AMPA as well as a combined AMPA and NMDA current inputs.

### 3.4 NMDA, Ca, and the K(Ca) current underlie nonmonotonic spike-adding

To explore how the relative amplitudes of AMPA and NMDA current input affect ADPs and spike-adding the conductances of the AMPA and NMDA currents were each varied. A nonmonotonic relationship between increasing *g*_*NMDA*_ and the number of evoked APs was found; indeed, a decrease in the number of evoked APs emerges as *g*_*NMDA*_ increases to 10 µS/cm^2^ (Figure 6a). This peculiar nonmonotonic spike-adding with increasing NMDA conductance can be modulated by the fraction of NMDA current that is carried by Ca^2+^ (*P*_*f,NMDA*_; Figure 6b) suggesting a Ca^2+^ dependent mechanism. Furthermore, scaling of the K(Ca) conductance can also modulate this nonmonotonic spike-adding (Figure 6c), further suggesting that the phenomenon is Ca^2+^ dependent and that it occurs through K(Ca) current. Modulation of the non-monotonic spike-adding and the ADPs associated with it as a result of alterations of the Hill coefficient and Ca^2+^ concentration at half activation (*k*_*Ca*_) of the Ca^2+^ -dependent gating of the K(Ca) current (Figure 6d and e respectively) provides additional support for a Ca^2+^ -dependent mechanism that activates a K(Ca) current, which in turn results in a nonmonotonic spike-adding with increasing NMDA conductance.

**Fig. 6.**
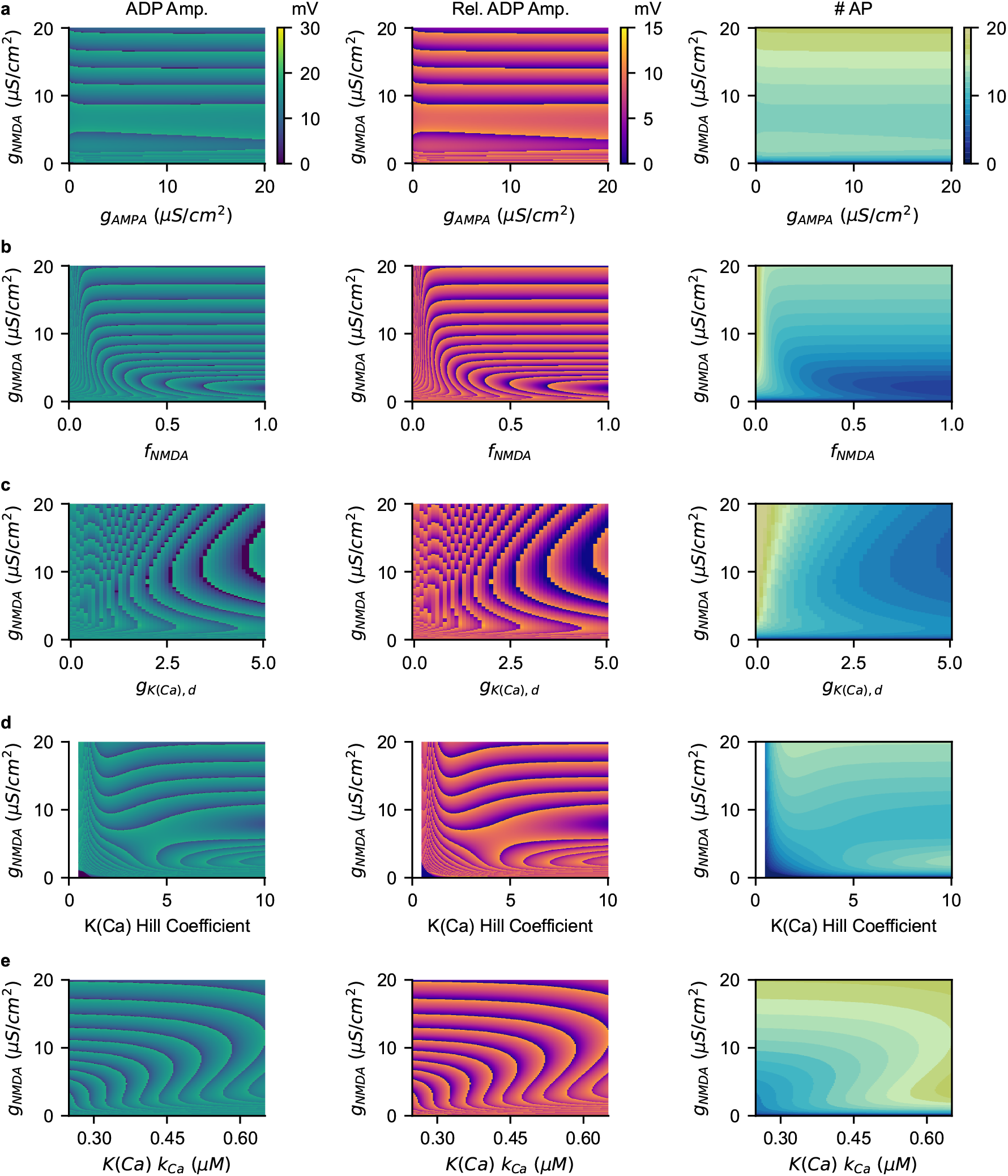
NMDA mediated Ca^2+^ effects on afterdepolarizations. The effect of varying the (a) conductances of AMPA and NMDA current input (a; *g*_*AMP A*_ and *g*_*NMDA*_, respectively), (b) conductance of NMDA current and the fraction of Ca^2+^ current carried by the NMDA current (*P*_*f,NMDA*_), (c) conductance of NMDA current and the K(Ca) current conductance (fold change from its default value) in the dendritic compartment (*g*_*K*(*Ca*),*d*_), (d) conductance of NMDA current and the the Hill coefficient of the gating of the K(Ca) current (*n*_*K*(*Ca*)_), and (d) conductance of NMDA current and the half-maximal concentration of the K(Ca) current gating (*k*_*Ca*_) on ADP amplitude, relative ADP amplitude and the number of evoked APs. A fixed AMPA conductance (*g*_*AMP A*_ = 10 µS/cm^2^) is assumed in b-e.

To probe such a mechanism, the integral of dendritic Ca^2+^ concentration ([*Ca*_*d*_]) and the K(Ca) current from the onset of the synaptic input to the last evoked AP were calculated and normalized by the number of evoked APs (Figure 7a-c) across increasing values of *g*_*NMDA*_. For low *g*_*NMDA*_ (left of the gray region, Figure 7d), the increase in the number of evoked APs is accompanied by a monotonic rise in ADP amplitudes up until a regime where the local maximum number (n=13) of APs is reached; in this latter regime, ADP amplitudes first rise and then decay with increasing *g*_*NMDA*_ (left of the gray region, Figure 7e). Beyond this regime at higher *g*_*NMDA*_, the number of APs decreases from the previous regime, while ADP amplitude initially decreases and then increases (gray region, Figure 7d and e, respectively) until another spike is added. Following this gray region, monotonic increases in ADP occur terminating in a spike-adding event as *g*_*NMDA*_ increases (right of the gray region, Figure 7d and e, respectively). The integral of both [*Ca*_*d*_] (Figure 7f) and K(Ca) (Figure 7g) across *g*_*NMDA*_ is similar to that of the number of evoked APs with a decrease in amplitude occurring during the decrease in APs evoked (in gray). However, when the rate of change in the ADP amplitude (Figure 7h), in the integral [*Ca*_*d*_] (Figure 7i) and in the integral of K(Ca) current (Figure 7j) were computed across each segment of consecutive *g*_*NMDA*_ that elicits the same number of AP, we found that the rate of change in ADP amplitude is minimal, the rate of change in integrated Ca^2+^ is increasing, and the rate of change of integrated K(Ca) current is maximal during the regime where the number of spikes is decreased (in gray, Figure 7h-j). As such, at low *g*_*NMDA*_ synaptic input induces spike-adding and multiple APs, thereby increasing [*Ca*_*d*_] and leading to a positive feedback loop that increases the number of evoked APs (Figure 7k). At a certain level of *g*_*NMDA*_, the increase in Ca^2+^ evoked by the APs is sufficiently large and activates the K(Ca) current, which in turn acts as negative feedback on the number of spikes evoked. If this negative feedback is sufficiently large, the number of APs decreases as *g*_*NMDA*_ increases (Figure 7l) resulting in the nonmonotonic spike-adding (highlighted in gray, Figure 7d-j). However, as *g*_*NMDA*_ increases further, the balance of positive and negative feedback from the elevation of [*Ca*_*d*_] reverts back to a net positive feedback (Figure 7m) and the number of evoked APs continues to increase. This is evident in the progressive growth of [*Ca*_*d*_] traces with increasing *g*_*NMDA*_ (middle of Figure 7k-m), whereas the amplitude of K(Ca) current saturates with larger NMDA conductance (bottom of Figure 7k-m).

**Fig. 7.**
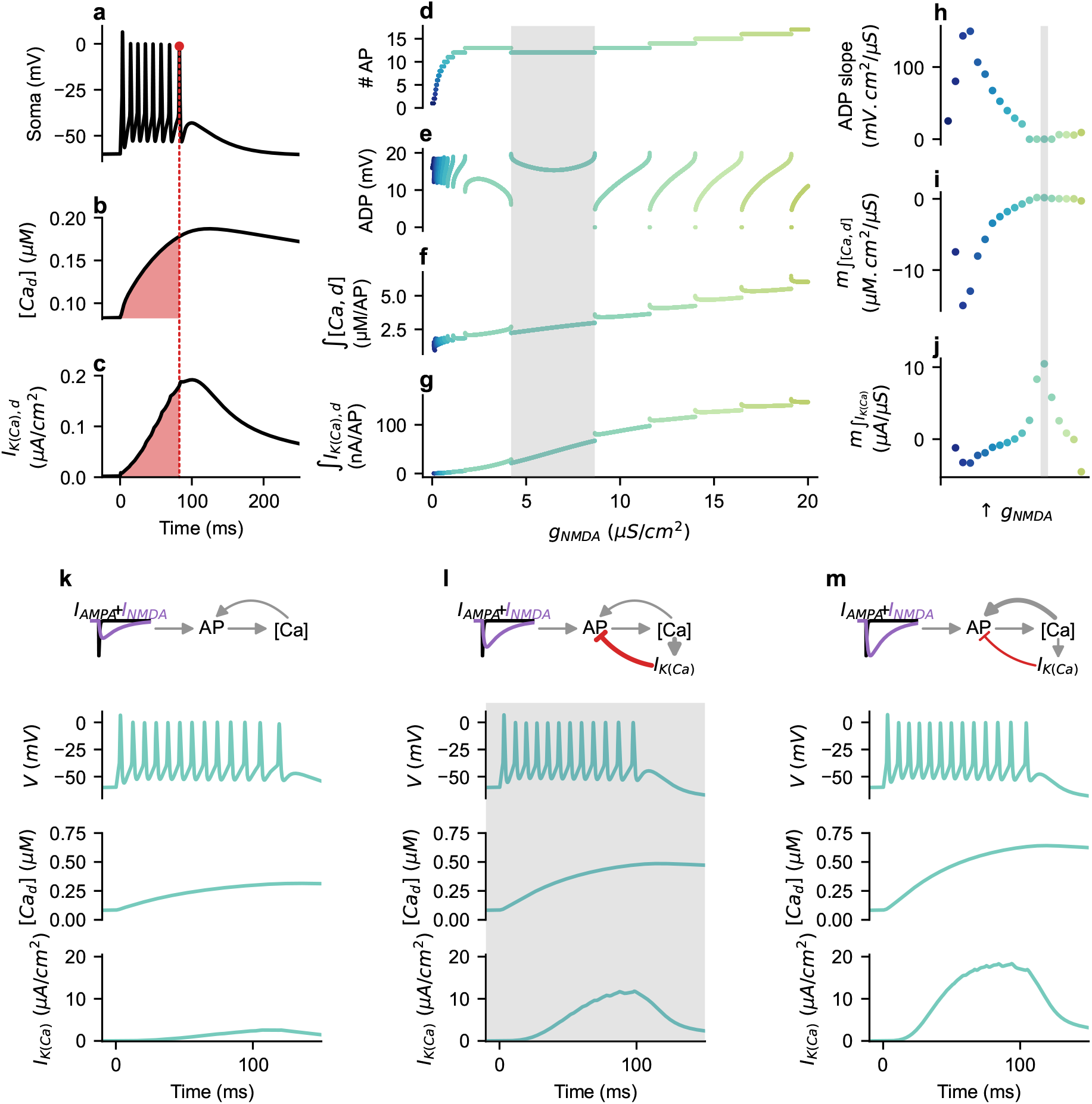
NMDA mediated spike-adding in the presence of Ca^2+^ and Ca^2+^ activated K^+^ currents. (a) An example somatic membrane potential response to an AMPA and NMDA current input. The last action potential is identified with a red dot, and the time at which it occurs is delineated by a dashed red line. (b) The integral of dendritic Ca^2+^ concentration ([*Ca*_*d*_]), and (c) dendritic K(Ca) current amplitude (*I*_*K*(*Ca*),*d*_) computed from the start of the input to the time of the last AP (red shading) and normalized by the difference of the input start time and the time of the last AP. (d) The number of APs elicited, (e) the amplitude of the ADP, and (f) the normalized integrals of [*Ca*_*d*_] and (g) *I*_*K*(*Ca*),*d*_ with increasing NMDA conductance (*g*_*NMDA*_). The range of NMDA conductance corresponding to a decrease in the number of evoked APs despite the increase in NMDA conductance is highlighted in gray. For each range of NMDA conductance that consistently evokes the same number of APs in d, the slope of the ADP amplitude (h), normalized dendritic Ca^2+^ (i) and dendritic K(Ca) current integral (j) with respect to NMDA conductance are computed. The gray region in h-j corresponds to the NMDA conductance range in gray in d-g. Example traces for *g*_*NMDA*_ = 2.5 µS/cm^2^ (k), 6.5 µS/cm^2^ (l), 10 µS/cm^2^ (m) with schematic of the effects of NMDA current input of AP generation, positive feedback of Ca^2+^ concentration ([Ca]) and negative feedback through K(Ca) current (*I*_*K*(*Ca*)_). The somatic membrane potential responses (top), dendritic Ca^2+^ concentration ([*Ca*_*d*_]; middle) and the dendritic K(Ca) current (*I*_*K*(*Ca*)_; bottom) are shown below each schematic. The example in l is in the gray region in d-g and h-j and is thus also shaded gray.

## 4 Discussion

Afterdepolarization potentials (ADPs) are the result of slow inward and fast outwards currents (Nowacki et al. 2013) and are prevalent throughout the nervous system, allowing them to modulate neuronal excitability (Gray and McCormick 1996; Amir et al. 2002; Chapin and Andrade 2000; Chen and Yaari 2008; Yue and Yaari 2006; D’Ascenzo et al. 2009; Connors et al. 1982; Greene et al. 1994; Staff et al. 2000; Li and Hatton 1997; Hilaire et al. 2005; Grampp 1966; Mayer 1985; Lemon and Turner 2000; Sheasby and Fohlmeister 1999; Stern and Armstrong 1996; Chu and Moenter 2006; Norekian 1999). Using a two compartmental model, a dendritic slow inward and fast outward currents were found to modulate ADP amplitudes and spike-adding to a larger degree compared to somatically located slow inward and fast outward currents. Dendritic morphology was also found to modulate ADPs and stimulus evoked transient bursting, with increasing dendritic size increasing ADPs and spike-adding, whereas increased dendritic leak conductance decreasing ADPs and spike-adding. Additionally, a nonmonotonic spike-adding pattern emerges with increasing NMDA current conductance, driven by the opposing effects of rising dendritic Ca^2+^ concentration and elevation in dendritic K(Ca) current.

The role of K(Ca) currents in the modulation of ADP amplitude and spike-adding is consistent with the roles of fast inward and slow outward currents on these phenomenon seen by Nowacki et al. (2013). However, the use of a two compartmental model in this study furthers these findings to demonstrate that the dendritic localization of these currents can modulate ADPs and spike-adding to a greater extent than somatically located currents. Additionally, in this study, we implemented a more realistic synaptic input time course (including AMPA and NMDA currents) to investigate how they mediate ADP generation and spike-adding rather than using brief step current inputs. Interestingly, our results revealed that with the slower kinetics of the NMDA current, stimulus-evoked transient bursts occur with many APs and that the number of APs evoked is modulated by fast inward and slow outward currents. In particular, a nonmonotonic spike-adding was found to occur as a result of the interactions of the slow NMDA current and slow Ca^2+^ timescales that activate the K(Ca) current. The role of such slow positive and negative feedback in providing this nonmonotonic spike-adding suggests dynamic regulation of stimulus evoked bursting by dendritic currents and highlights that interactions of processes of similar timescales (slower than AP generation) may modulate ADPs and spike-adding.

### 4.1 Dendritic size modulates ADPs and stimulus evoked transient bursting in a mechanism-dependent manner

In the two compartmental model, larger dendrites or increases in 1 ™ *κ* result in increased ADPs in response to both dendritic step current inputs and AMPA current inputs as well as increased number of evoked APs by NMDA current input (Figure 3a-c). This is in direct contradiction to the decreases in ADPs seen with increasing dendritic length in retinal ganglion cell (Sheasby and Fohlmeis- ter 1999), hypothalamic magnocellular neurosecretory cells (Stern and Armstrong 1996, 1998; Teruyama and Armstrong 2002), gonadotropin-releasing hormone (GnRH) neurons (Roberts et al. 2008), but are in line with the decreases in ADPs seen with decreased dendritic size in two compartmental models (Mainen and Sejnowski 1996). It should be noted, however, that these findings in retinal ganglion cell, magnocellular neurosecretory cells and GnRH neurons were obtained through somatic current injection during patchclamp recording, and not with dendritic current inputs as in Figure 3. Additionally, ADPs and bursting were elucidated through extracellular ion concentration changes and pharmacological means in pyramidal neurons with truncated dendrites (Golomb et al. 2006), suggesting that the mechanism underlying ADPs and the stimulus-evoked transient bursting in these neurons is likely somatic and not dependent on dendrites.

With this in mind, it seems likely that neurons with exclusively somatic ADP generating mechanisms display reductions in ADP amplitude and stimulus evoked transient bursting as dendritic size increases due to the shunting effect of the dendritic leak conductance; in contrast, neurons where ADPs are generated by dendrite-dependent mechanisms (e.g. by elevation in dendritic Ca^2+^ concentration) exhibit increases in ADPs and transient bursting with increasing dendritic size because of increased compartmentalization of distal dendrites from the soma (i.e. dendritic voltage and Ca^2+^ fluctuations are less tied to somatic ones). In line with this hypothesis, two compartmental models of pyramidal neurons that are reliant on the interaction between fast somatic conductances and slow dendritic conductances for the generation of ADPs have larger ADPs with increased dendritic size (Mainen and Sejnowski 1996). Additionally, in the two compartmental model presented here where dendritic mechanisms contribute to ADP generation, the effect of increasing the dendritic size (i.e. 1 ™ *κ*) with somatic instead of dendritic step current inputs results in modest increases in ADP amplitude (Figure S5a-c) and increases in dendritic leak modestly decreasing ADP amplitudes (Figure S5d-f), suggesting no qualitative input location dependence in the nature of the relationship between dendritic length or dendritic leak conductance with ADPs. Furthermore, the reduction in ADPs with increased dendritic leak conductance (Figure 3 and S2) corresponds to the proposed shunting role of the passive (i.e. leak) properties of dendrites on ADPs (Roberts et al. 2008). However, intermediate range of dendritic size and somato-dendritic coupling has been shown to be required for stimulus-driven transient bursting (Pinsky and Rinzel 1994; Yi et al. 2025; van Elburg and van Ooyen 2010), in line with the range of dendritic sizes that support ADPs in the model with cable dendrite (Figure S2f), but not the two compartmental model (Figure S2c), suggesting a limitation in the approximation of the dendritic arbor into a single compartment.

Dendritic compartmentalization and integration are known to contribute to neuronal computations (Francioni and Harnett 2022), indicating that dendritic processes contributing to ADPs and stimulus evoked transient bursting may provide computational advantages. Indeed, backpropagating APs in CA1 pyramidal neurons can result in bursts of firing when they encounter depolarized dendrites, in a manner dependent on A-type K^+^ channels (Park et al. 2025). Specifically, the inactivation of A-type K^+^ channels and the Na^+^ channel slow inactivation define a window in time such that dendritic spikes only occur with somatic stimulation if the dendrites are depolarized during this window (Park et al. 2025). However, back-propagating APs can also interact with stimulus evoked dendritic depolarization to inactivate A-type K^+^ channels, enabling dendritic spiking and stimulus evoked transient bursting (Park et al. 2025), in a manner similar to the role of this current in modulating ADPs. Additionally, ADPs are not passively propagated along dendrites but filtered (Kalmbach et al. 2013), making it likely that the dendritic currents and mechanisms involved in regulating ADPs and spike-adding contribute to dendritic computations to regulate neuronal excitability and plasticity.

### 4.2 Cerebellar stellate cells with extrasynaptic NMDA receptors have ADPs and synaptically driven transient bursting

Cerebellar stellate cells (CSCs) are GABAergic molecular layer interneurons in the molecular layer of the cerebellum that inhibit Purkinje cells (Liu et al. 2014) and whose excitability can be regulated by shifts in the voltage dependence of Na^+^ and A-type K^+^ channels (Alexander et al. 2019). CSCs have Ca^2+^ permeable AMPA receptors (Tran-Van-Minh et al. 2016; Rossi et al. 2008) and extrasynaptic GluN2B containing NMDA receptors (Carter and Regehr 2000; Nahir and Jahr 2013; Bidoret et al. 2015). High frequency parallel fiber (Nahir and Jahr 2013; Carter and Regehr 2000; Clark and Cull-Candy 2002) or concurrent parallel and climbing fiber activity (Szapiro and Barbour 2007; Jörntell and Ekerot 2003, 2002) results in glutamate spillover that activates the extrasynaptic NMDA receptors to evoke ADPs and transient bursts of APs in a frequency dependent manner (Carter and Regehr 2000; Clark and Cull-Candy 2002) which contribute to irregularity in *in vivo* CSC firing patterns (Suter and Jaeger 2004; Jörntell et al. 2010). These NMDA receptor evoked increases in molecular layer interneuron excitability result in rapid and strong inhibition of Purkinje cells to modulate cerebellar cortex information processing (Liu et al. 2014). Additionally, NMDA receptor activation in CSCs is likely to control synaptic plasticity, with NMDA receptors required for dendritic Ca^2+^ evoked short term depression (Tran-Van-Minh et al. 2016) and for nitric oxide dependent long term depression (Kono et al. 2018). Knockout of molecular layer interneuron NMDA receptors results in impairment in a cerebellar learning task (Kono et al. 2018) highlighting the importance of such mechanisms in behavior. Additionally, ADPs play a role during development in CSCs, with *GABA*_*A*_ autoreceptor activation resulting in ADPs and transient bursting, providing a a mechanism to facilitate the stabilization of inhibitory synapses during the developmental period when inhibitory synapses are stabilizing (Mejia-Gervacio and Marty 2006).

### 4.3 ADP generating mechanisms and nonmonotonic spike-adding

Many different slow currents, such as persistent Na^+^ (Mainen and Sejnowski 1996; Chen and Yaari 2008; Park et al. 2025), h-current (*I*_*h*_) (Chapin and Andrade 2000; Kalmbach et al. 2013), Ca^2+^ activated non-selective cationic (CAN) currents (Shpak et al. 2015; Hasuo et al. 1990), as well as the removal of K^+^ currents (Greene et al. 1994; Sultan et al. 2024; Li and Hatton 1997), have been shown to underlie ADPs and spike-adding. The NMDA-dependent nonmonotonic spike-adding effect described here is likely not dependent on the mechanism by which ADPs and spike-adding occurs, but rather by the occurrence of ADPs and the slower timescales of the NMDA evoked Ca^2+^ influx and resulting K(Ca) current in the dendritic compartment of a neuron with dendritic-dependent ADP generation mechanism. As such, any mechanisms that result in oppositional currents of similar timescales is likely to generate this phenomenon. For example, neuromodulators, such as noradrenergic (Yoshimura et al. 1987), D2 dopaminergic (Evans et al. 2017), and mGluR5 receptor (D’Ascenzo et al. 2009) activation can increase ADPs and neuronal excitability, in some cases only in neuronal subpopulations (Evans et al. 2017). As such, the slower timescale of neuromodulators presents the possibility that nonmonotonic increases in spike-adding, describe here with NMDA current input, may exist as the result of interactions between the effects of neuromodulators and the slow outward currents underlying ADPs. Additionally, these mechanisms and the tight regulation of spike-adding and stimulus evoked transient bursting through inward and outward currents and the dendritic Ca^2+^ influx they evoked are likely coupled with mechanisms of synaptic plasticity (Dudai et al. 2021; Park et al. 2025), although further work is required to substantiate this.

### 4.4 Limitations

The approximation of the entire dendritic tree into a single dendritic compartment, as is the case for the two compartmental model, neglects the spatial aspect and topology of the dendrites. While this is likely not an issue for the qualitative results, some discrepancies, such as those noted with the absence of a range of dendritic lengths supporting ADPs and transient bursting in the two compartmental model, may occur as a result. Additionally, at sufficiently small coupling, the dendritic compartment is electrotonically isolated from the soma in a manner similar to some distal dendrites. However, any active filtering that occurs in more proximal dendrites between these distal dendrites and the soma is neglected. Moreover, other mechanisms of ADP generation mentioned above such as persistent Na^+^ currents (Mainen and Sejnowski 1996; Chen and Yaari 2008; Park et al. 2025), *I*_*h*_ (Chapin and Andrade 2000; Kalmbach et al. 2013), Ca^2+^ activated nonselective cation currents (Shpak et al. 2015; Hasuo et al. 1990), and removal of K^+^ currents (Greene et al. 1994; Sultan et al. 2024; Li and Hatton 1997) were not explicitly explored. However, as the nonmonotonic spike-adding occurs as a direct consequence of the interaction between the NMDA current and the slow Ca^2+^ and K(Ca) current, it is likely that these other mechanisms could also result in nonmonotonic spike-adding if dendritic Ca^2+^ is increased or slow depolarizing and hyperpolarizing currents are activated in a similar manner. Finally, all simulations performed in this study were done in the absence of background synaptic activity and as such how these behaviors translate to those conditions is not known, but increased variability in ADPs and stimulus evoked bursting are expected.

## Declarations

### Funding

This work was supported by the Natural Sciences and Engineering Research Council of Canada (NSERC) Discovery (RGPIN-2019-04520) and Alliance International (ALLRP 588367-23) grants to AK. NK was supported by the Canada Graduate Research Scholarship – Doctoral program. whereas YZ was supported by Health Brains, Health Lives as well as the Mackey-Glass fellowships. This research was enabled in part by support provided by Calcul Québec (https://www.calculquebec.ca) and the Digital Research Alliance of Canada (https://alliancecan.ca/). The funders had no role in study design, data collection and analysis, decision to publish, or preparation of the manuscript.

### Competing Interests

The authors declare no competing interests.

### Data availability

No datasets were generated or analysed during the current study.

### Code availability

The code used to generate the data and figures are available at https://github.com/nkoch1/ADP Koch et al.git.

### Author Contribution

**N.A.K**: Conceptualization, Methodology, Software, Validation, Formal analysis, Investigation, Writing - Original Draft, Writing - Review & Editing, Visualization. **Y.Z**.: Methodology, Formal analysis, Investigation, Writing - Review & Editing **A.K**.: Conceptualization, Resources, Writing - Review & Editing, Supervision, Funding acquisition.

## Notes

### Competing Interest Statement

The authors have declared no competing interest.

## References

Abrahamsson T, Cathala L, Matsui K, Shigemoto R, DiGregorio DA. Thin Dendrites of Cerebellar Interneurons Confer Sublinear Synaptic Integration and a Gradient of Short-Term Plasticity. Neuron. 2012;73(6):1159–1172. 10.1016/j.neuron.2012.01.027.

Akhshi A, Haggard M, Marquez MM, Farjami S, Chacron MJ, Khadra A. Decoding the relative contributions of extrinsic and intrinsic mechanisms in mediating heterogeneous spiking activities of sensory neurons in vivo using computational modeling. bioRxiv. 2023 Jan;10.1101/2023.01.03.521866.

Akhshi A, Metzen MG, Chacron MJ, Khadra A. In vivo neural activity of electrosensory pyramidal cells: Biophysical characterization and phenomenological modeling. bioRxiv. 2025 Jun;10.1101/2025.05.30.656684.

Alexander RPD, Mitry J, Sareen V, Khadra A, Bowie D. Cerebellar Stellate Cell Excitability Is Coordinated by Shifts in the Gating Behavior of Voltage-Gated Na+ and A-Type K+ Channels. eNeuro. 2019;6(3):ENEURO.0126–19.2019. 10.1523/ENEURO.0126-19.2019.

Amir R, Michaelis M, Devor M. Burst Discharge in Primary Sensory Neurons: Triggered by Subthreshold Oscillations, Maintained by Depolarizing Afterpotentials. The Journal of Neuroscience. 2002;22(3):1187–1198. 10.1523/jneurosci.22-03-01187.2002.

Beaver ML, Evans RC. Muscarinic Receptor Activation Preferentially Inhibits Rebound in Vulnerable Dopaminergic Neurons. The Journal of Neuroscience. 2025;45(16):e1443242025. 10.1523/jneurosci.1443-24.2025.

Bezanson J, Edelman A, Karpinski S, Shah VB. Julia: A Fresh Approach to Numerical Computing. SIAM Rev. 2017;59(1):65–98. 10.1137/141000671.

Bidoret C, Bouvier G, Ayon A, Szapiro G, Casado M. Properties and molecular identity of NMDA receptors at synaptic and non-synaptic inputs in cerebellar molecular layer interneurons. Frontiers in Synaptic Neuroscience. 2015;7. 10.3389/fnsyn.2015.00001.

Brunel N, Wang XJ. Effects of Neuromodulation in a Cortical Network Model of Object Working Memory Dominated by Recurrent Inhibition. Journal of Computational Neuroscience. 2001;11(1):63–85. 10.1023/a:1011204814320.

Burnashev N, Zhou Z, Neher E, Sakmann B. Fractional calcium currents through recombinant GluR channels of the NMDA, AMPA and kainate receptor subtypes. The Journal of Physiology. 1995;485(2):403–418. 10.1113/jphysiol.1995.sp020738.

Burnashev N. Calcium permeability of glutamate-gated channels in the central nervous system. Current Opinion in Neurobiology. 1996;6(3):311–317. 10.1016/s0959-4388(96)80113-9.

Carannante I, Johansson Y, Silberberg G, Hellgren Kotaleski J. Data-Driven Model of Postsynaptic Currents Mediated by NMDA or AMPA Receptors in Striatal Neurons. Frontiers in Computational Neuroscience. 2022;16. 10.3389/fncom.2022.806086.

Carter AG, Regehr WG. Prolonged Synaptic Currents and Glutamate Spillover at the Parallel Fiber to Stellate Cell Synapse. The Journal of Neuroscience. 2000;20(12):4423–4434. 10.1523/jneurosci.20-12-04423.2000.

Chapin EM, Andrade R. Calcium-Independent Afterdepolarization Regulated by Serotonin in Anterior Thalamus. Journal of Neurophysiology. 2000;83(5):3173–3176. 10.1152/jn.2000.83.5.3173.

Chen S, Yaari Y. Spike Ca2+influx upmodulates the spike afterdepolarization and bursting via intracellular inhibition of KV7/M channels. The Journal of Physiology. 2008;586(5):1351–1363. 10.1113/jphysiol.2007.148171.

Chu Z, Moenter SM. Physiologic Regulation of a Tetrodotoxin-Sensitive Sodium Influx That Mediates a Slow Afterdepolarization Potential in Gonadotropin-Releasing Hormone Neurons: Possible Implications for the Central Regulation of Fertility. The Journal of Neuroscience. 2006;26(46):11961–11973. 10.1523/jneurosci.3171-06.2006.

Clark BA, Cull-Candy SG. Activity-Dependent Recruitment of Extrasynaptic NMDA Receptor Activation at an AMPA Receptor-Only Synapse. The Journal of Neuroscience. 2002;22(11):4428–4436. 10.1523/jneurosci.22-11-04428.2002.

Connors BW, Gutnick MJ, Prince DA. Electrophysiological properties of neocortical neurons in vitro. Journal of Neurophysiology. 1982;48(6):1302–1320. 10.1152/jn.1982.48.6.1302.

Dayan P, Abbott L, Dayan P, Abbott LF. Theoretical Neuroscience: Computational and Mathematical Modeling of Neural Systems. Computational neuroscience, MIT Press; 2001.

Dixit VK, Rackauckas C. GlobalSensitivity.jl: Performant and Parallel Global Sensitivity Analysis with Julia. Journal of Open Source Software. 2022;7(76):4561.

Dixit VK, Rackauckas C.: Optimization.jl: A Unified Optimization Package. Zenodo; 2023. 10.5281/zenodo.7738525.

Doiron B, Laing C, Longtin A, Maler L. Ghost-bursting: A Novel Neuronal Burst Mechanism. Journal of Computational Neuroscience. 2002;12(1):5–25. 10.1023/a:1014921628797.

Doiron B, Noonan L, Lemon N, Turner RW. Persistent Na+Current Modifies Burst Discharge By Regulating Conditional Backpropagation of Dendritic Spikes. Journal of Neurophysiology. 2003;89(1):324–337. 10.1152/jn.00729.2002.

Dudai A, Doron M, Segev I, London M. Synaptic Input and ACh Modulation Regulate Dendritic Ca2+Spike Duration in Pyramidal Neurons, Directly Affecting Their Somatic Output. The Journal of Neuroscience. 2021;42(7):1184–1195. 10.1523/jneurosci.1470-21.2021.

D’Ascenzo M, Podda MV, Fellin T, Azzena GB, Haydon P, Grassi C. Activation of mGluR5 induces spike afterdepolarization and enhanced excitability in medium spiny neurons of the nucleus accumbens by modulating persistent Na+ currents. The Journal of Physiology. 2009;587(13):3233–3250. 10.1113/jphysiol.2009.172593.

van Elburg RAJ, van Ooyen A. Impact of Dendritic Size and Dendritic Topology on Burst Firing in Pyramidal Cells. PLoS Computational Biology. 2010;6(5):e1000781. 10.1371/journal.pcbi.1000781.

Evans RC, Zhu M, Khaliq ZM. Dopamine Inhibition Differentially Controls Excitability of Substantia Nigra Dopamine Neuron Subpopulations through T-Type Calcium Channels. The Journal of Neuroscience. 2017;37(13):3704–3720. 10.1523/jneurosci.0117-17.2017.

Farjami S, Alexander RPD, Bowie D, Khadra A. Bursting in cerebellar stellate cells induced by pharmacological agents: Non-sequential spike adding. PLOS Computational Biology. 2020;16(12):e1008463. 10.1371/journal.pcbi.1008463.

Farjami S, Alexander RPD, Bowie D, Khadra A. Switching in Cerebellar Stellate Cell Excitability in Response to a Pair of Inhibitory/Excitatory Presynaptic Inputs: A Dynamical System Perspective. Neural Computation. 2020;32(3):626–658. 10.1162/neco_a_01261.

Francioni V, Harnett MT. Rethinking Single Neuron Electrical Compartmentalization: Dendritic Contributions to Network Computation In Vivo. Neuroscience. 2022;489:185–199. 10.1016/j.neuroscience.2021.05.038.

Fransén E, Alonso AA, Hasselmo ME. Simulations of the Role of the Muscarinic-Activated Calcium-Sensitive Nonspecific Cation Current INCM in Entorhinal Neuronal Activity during Delayed Matching Tasks. The Journal of Neuroscience. 2002;22(3):1081–1097. 10.1523/jneurosci.22-03-01081.2002.

Gao T, Deng B, Wang J, Wang J, Yi G. The passive properties of dendrites modulate the propagation of slowly-varying firing rate in feedforward networks. Neural Networks. 2022;150:377–391. 10.1016/j.neunet.2022.03.001.

Golomb D, Yue C, Yaari Y. Contribution of Persistent Na+Current and M-Type K+Current to Somatic Bursting in CA1 Pyramidal Cells: Combined Experimental and Modeling Study. Journal of Neurophysiology. 2006;96(4):1912–1926. 10.1152/jn.00205.2006.

Grampp W. Multiple-Spike Discharge Evoking After-Depolarizations in the Slowly Adapting Stretch Receptor Neuron of the Lobster II. The slow after-depolarization. Acta Physiologica Scandinavica. 1966;67(1):116–126. 10.1111/j.1748-1716.1966.tb03292.x.

Gray CM, McCormick DA. Chattering Cells: Superficial Pyramidal Neurons Contributing to the Generation of Synchronous Oscillations in the Visual Cortex. Science. 1996;274(5284):109–113. 10.1126/science.274.5284.109.

Greene CC, Schwindt PC, Crill WE. Properties and ionic mechanisms of a metabotropic glutamate receptor-mediated slow afterdepolarization in neocortical neurons. Journal of Neurophysiology. 1994;72(2):693–704. 10.1152/jn.1994.72.2.693.

Gu N, Vervaeke K, Hu H, Storm JF. Kv7/KCNQ/M and HCN/h, but not KCa2/SK channels, contribute to the somatic medium after-hyperpolarization and excitability control in CA1 hippocampal pyramidal cells. The Journal of Physiology. 2005;566(3):689–715. 10.1113/jphysiol.2005.086835.

Hasuo H, Phelan KD, Twery MJ, Gallagher JP. A calcium-dependent slow afterdepolarization recorded in rat dorsolateral septal nucleus neurons in vitro. Journal of Neurophysiology. 1990;64(6):1838–1846. 10.1152/jn.1990.64.6.1838.

Higashi H, Tanaka E, Inokuchi H, Nishi S. Ionic mechanisms underlying the depolarizing and hyperpolarizing afterpotentials of single spike in guinea-pig cingulate cortical neurons. Neuroscience. 1993;55(1):129–138. 10.1016/0306-4522(93)90460-w.

Hilaire C, Campo B, André S, Valmier J, Scamps F. K+ current regulates calcium-activated chloride current-induced afterdepolarization in axotomized sensory neurons. European Journal of Neuroscience. 2005;22(5):1073–1080. 10.1111/j.1460-9568.2005.04271.x.

Jahr C, Stevens C. Voltage dependence of NMDA-activated macroscopic conductances predicted by single-channel kinetics. The Journal of Neuroscience. 1990;10(9):3178–3182. 10.1523/jneurosci.10-09-03178.1990.

Jansen MJW. Analysis of variance designs for model output. Computer Physics Communications. 1999;117(1–2):35–43. 10.1016/s0010-4655(98)00154-4.

Johnson SG.: The NLopt nonlinear-optimization package; 2007. https://github.com/stevengj/nlopt.

Jung Hy, Staff NP, Spruston N. Action Potential Bursting in Subicular Pyramidal Neurons Is Driven by a Calcium Tail Current. The Journal of Neuroscience. 2001;21(10):3312–3321. 10.1523/jneurosci.21-10-03312.2001.

Jörntell H, Bengtsson F, Schonewille M, De Zeeuw CI. Cerebellar molecular layer interneurons – computational properties and roles in learning. Trends in Neurosciences. 2010;33(11):524–532. 10.1016/j.tins.2010.08.004.

Jörntell H, Ekerot CF. Reciprocal Bidirectional Plasticity of Parallel Fiber Receptive Fields in Cerebellar Purkinje Cells and Their Afferent Interneurons. Neuron. 2002;34(5):797–806. 10.1016/s0896-6273(02)00713-4.

Jörntell H, Ekerot CF. Receptive Field Plasticity Profoundly Alters the Cutaneous Parallel Fiber Synaptic Input to Cerebellar InterneuronsIn Vivo. The Journal of Neuroscience. 2003;23(29):9620–9631. 10.1523/jneurosci.23-29-09620.2003.

Kalmbach BE, Chitwood RA, Dembrow NC, Johnston D. Dendritic Generation of mGluR-Mediated Slow Afterdepolarization in Layer 5 Neurons of Prefrontal Cortex. The Journal of Neuroscience. 2013;33(33):13518–13532. 10.1523/jneurosci.2018-13.2013.

Kleppe IC, Robinson HPC. Determining the Activation Time Course of Synaptic AMPA Receptors from Openings of Colocalized NMDA Receptors. Biophysical Journal. 1999;77(3):1418–1427. 10.1016/s0006-3495(99)76990-0.

Klinshov VV, Nekorkin VI. Activity clusters in dynamical model of the working memory system. Network: Computation in Neural Systems. 2008;19(2):119–135. 10.1080/09548980801895452.

Klinshov VV, Nekorkin VI. Delayed afterdepolarization and spontaneous secondary spiking in a simple model of neural activity. Communications in Nonlinear Science and Numerical Simulation. 2012;17(3):1438–1446. 10.1016/j.cnsns.2011.08.020.

Kono M, Kakegawa W, Yoshida K, Yuzaki M. Interneuronal NMDA receptors regulate long-term depression and motor learning in the cerebellum. The Journal of Physiology. 2018;597(3):903–920. 10.1113/jp276794.

Krichmar JL, Nasuto SJ, Scorcioni R, Washington SD, Ascoli GA. Effects of dendritic morphology on CA3 pyramidal cell electrophysiology: a simulation study. Brain Research. 2002;941(1–2):11–28. 10.1016/s0006-8993(02)02488-5.

Lehnert J, Khadra A. How Pulsatile Kisspeptin Stimulation and GnRH Autocrine Feedback Can Drive GnRH Secretion: A Modeling Investigation. Endocrinology. 2019 Mar;160(5):1289–1306. 10.1210/en.2018-00947.

Lemon N, Turner RW. Conditional Spike Back-propagation Generates Burst Discharge in a Sensory Neuron. Journal of Neurophysiology. 2000;84(3):1519–1530. 10.1152/jn.2000.84.3.1519.

Li Z, Hatton GI. Reduced outward K+ conductances generate depolarizing after–potentials in rat supraoptic nucleus neurones. The Journal of Physiology. 1997;505(1):95–106. 10.1111/j.1469-7793.1997.095bc.x.

Lisman JE, Idiart MAP. Storage of 7 ± 2 Short-Term Memories in Oscillatory Subcycles. Science. 1995;267(5203):1512–1515. 10.1126/science.7878473.

Liu H, Zhao SN, Zhao GY, Sun L, Chu CP, Qiu DL. N-methyl-d-aspartate inhibits cerebellar Purkinje cell activity via the excitation of molecular layer interneurons under in vivo conditions in mice. Brain Research. 2014;1560:1–9. 10.1016/j.brainres.2014.03.011.

Liu X, Herbison AE. Small-Conductance Calcium-Activated Potassium Channels Control Excitability and Firing Dynamics in Gonadotropin-Releasing Hormone (GnRH) Neurons. Endocrinology. 2008;149(7):3598–3604. 10.1210/en.2007-1631.

Magee JC, Carruth M. Dendritic Voltage-Gated Ion Channels Regulate the Action Potential Firing Mode of Hippocampal CA1 Pyramidal Neurons. Journal of Neurophysiology. 1999;82(4):1895–1901. 10.1152/jn.1999.82.4.1895.

Mainen ZF, Sejnowski TJ. Influence of dendritic structure on firing pattern in model neocortical neurons. Nature. 1996;382(6589):363–366. 10.1038/382363a0.

Matoušek J. On theL2-Discrepancy for Anchored Boxes. Journal of Complexity. 1998;14(4):527–556. 10.1006/jcom.1998.0489.

Mayer ML. A calcium-activated chloride current generates the after-depolarization of rat sensory neurones in culture. The Journal of Physiology. 1985;364(1):217–239. 10.1113/jphysiol.1985.sp015740.

Mejia-Gervacio S, Marty A. Control of interneurone firing pattern by axonal autoreceptors in the juvenile rat cerebellum. The Journal of Physiology. 2006;571(Pt 1):43–55. 10.1113/jphysiol.2005.101675.

Metz AE. R-Type Calcium Channels Contribute to Afterdepolarization and Bursting in Hippocampal CA1 Pyramidal Neurons. Journal of Neuroscience. 2005;25(24):5763–5773. 10.1523/jneurosci.0624-05.2005.

Metz AE, Spruston N, Martina M. Dendritic D-type potassium currents inhibit the spike afterdepolarization in rat hippocampal CA1 pyramidal neurons. The Journal of Physiology. 2007;581(1):175–187. 10.1113/jphysiol.2006.127068.

Metzen MG, Akhshi A, Bashivan P, Khadra A, Chacron MJ. Burst firing optimizes invariant coding of natural communication signals by electrosensory neural populations. iScience. 2025 May;28(5):112399. 10.1016/j.isci.2025.112399.

Mogensen PK, Riseth AN. Optim: A mathematical optimization package for Julia. Journal of Open Source Software. 2018;3(24):615. 10.21105/joss.00615.

Moran S, Moenter SM, Khadra A. A unified model for two modes of bursting in GnRH neurons. Journal of Computational Neuroscience. 2016 Mar;40(3):297–315. 10.1007/s10827-016-0598-4.

Nahir B, Jahr CE. Activation of Extrasynaptic NMDARs at Individual Parallel Fiber–Molecular Layer Interneuron Synapses in Cerebellum. The Journal of Neuroscience. 2013;33(41):16323–16333. 10.1523/jneurosci.1971-13.2013.

Niday Z, Bean BP. BK Channel Regulation of Afterpotentials and Burst Firing in Cerebellar Purkinje Neurons. The Journal of Neuroscience. 2021;41(13):2854–2869. 10.1523/jneurosci.0192-20.2021.

Norekian TP. GABAergic Excitatory Synapses and Electrical Coupling Sustain Prolonged Discharges in the Prey Capture Neural Network ofClione limacina. The Journal of Neuroscience. 1999;19(5):1863–1875. 10.1523/jneurosci.19-05-01863.1999.

Nowacki J, Osinga HM, Brown JT, Randall AD, Tsaneva-Atanasova K. A unified model of CA1/3 pyramidal cells: An investigation into excitability. Progress in Biophysics and Molecular Biology. 2011;105(1–2):34–48. 10.1016/j.pbiomolbio.2010.09.020.

Nowacki J, Osinga HM, Tsaneva-Atanasova K. Dynamical systems analysis of spike-adding mechanisms in transient bursts. The Journal of Mathematical Neuroscience. 2012;2(1). 10.1186/2190-8567-2-7.

Nowacki J, Osinga HM, Tsaneva-Atanasova KT. Continuation-Based Numerical Detection of After-Depolarization and Spike-Adding Thresholds. Neural Computation. 2013;25(4):877–900. 10.1162/NECO_a_00425.

Okamoto H, Fukai T. Recurrent Network Models for Perfect Temporal Integration of Fluctuating Correlated Inputs. PLoS Computational Biology. 2009;5(6):e1000404. 10.1371/journal.pcbi.1000404.

van Ooyen A, Duijnhouwer J, Remme MWH, van Pelt J. The effect of dendritic topology on firing patterns in model neurons. Network: Computation in Neural Systems. 2002;13(3):311. 10.1088/0954-898X/13/3/304.

Park P, Wong-Campos JD, Itkis DG, Lee BH, Qi Y, Davis HC, et al. Dendritic excitations govern back-propagation via a spike-rate accelerometer. Nature Communications. 2025;16(1). 10.1038/s41467-025-55819-9.

Ping HX, Shepard PD. Blockade of SK-Type Ca2+-Activated K+Channels Uncovers a Ca2+-Dependent Slow Afterdepolarization in Nigral Dopamine Neurons. Journal of Neurophysiology. 1999;81(3):977–984. 10.1152/jn.1999.81.3.977.

Pinsky PF, Rinzel J. Intrinsic and network rhythmogenesis in a reduced traub model for CA3 neurons. Journal of Computational Neuroscience. 1994;1(1–2):39–60. 10.1007/bf00962717.

Poolos NP, Johnston D. Calcium-Activated Potassium Conductances Contribute to Action Potential Repolarization at the Soma But Not the Dendrites of Hippocampal CA1 Pyramidal Neurons. The Journal of Neuroscience. 1999;19(13):5205–5212. 10.1523/jneurosci.19-13-05205.1999.

Rackauckas C, Nie Q. DifferentialEquations.jl – A Performant and Feature-Rich Ecosystem for Solving Differential Equations in Julia. Journal of Open Research Software. 2017;10.5334/jors.151.

Rizza MF, Locatelli F, Masoli S, Sánchez-Ponce D, Muñoz A, Prestori F, et al. Stellate cell computational modeling predicts signal filtering in the molecular layer circuit of cerebellum. Scientific Reports. 2021;11(1):3873. 10.1038/s41598-021-83209-w.

Roberts CB, O’Boyle MP, Suter KJ. Dendrites determine the contribution of after depolarization potentials (ADPs) to generation of repetitive action potentials in hypothalamic gonadotropin releasing-hormone (GnRH) neurons. Journal of Computational Neuroscience. 2008;26(1):39–53. 10.1007/s10827-008-0095-5.

Rossi B, Maton G, Collin T. Calcium-permeable presynaptic AMPA receptors in cerebellar molecular layer interneurones. The Journal of Physiology. 2008;586(21):5129–5145. 10.1113/jphysiol.2008.159921.

Rowan TH. Functional stability analysis of numerical algorithms. PhD thesis, Department of Computer Science, University of Texas at Austin; 1990.

Saltelli A, Ratto M, Andres T, Campolongo F, Cariboni J, Gatelli D, et al. Global sensitivity analysis: the primer. John Wiley & Sons; 2008.

Sheasby BW, Fohlmeister JF. Impulse Encoding Across the Dendritic Morphologies of Retinal Ganglion Cells. Journal of Neurophysiology. 1999;81(4):1685–1698. 10.1152/jn.1999.81.4.1685.

Shpak G, Zylbertal A, Wagner S. Transient and sustained afterdepolarizations in accessory olfactory bulb mitral cells are mediated by distinct mechanisms that are differentially regulated by neuromodulators. Frontiers in Cellular Neuroscience. 2015;8. 10.3389/fncel.2014.00432.

Sobol IM. Global sensitivity indices for nonlinear mathematical models and their Monte Carlo estimates. Mathematics and Computers in Simulation. 2001;55(1–3):271–280. 10.1016/s0378-4754(00)00270-6.

Staff NP, Jung HY, Thiagarajan T, Yao M, Spruston N. Resting and Active Properties of Pyramidal Neurons in Subiculum and CA1 of Rat Hippocampus. Journal of Neurophysiology. 2000;84(5):2398–2408. 10.1152/jn.2000.84.5.2398.

Stern JE, Armstrong WE. Changes in the Electrical Properties of Supraoptic Nucleus Oxytocin and Vasopressin Neurons during Lactation. The Journal of Neuroscience. 1996;16(16):4861–4871. 10.1523/jneurosci.16-16-04861.1996.

Stern JE, Armstrong WE. Reorganization of the Dendritic Trees of Oxytocin and Vasopressin Neurons of the Rat Supraoptic Nucleus during Lactation. The Journal of Neuroscience. 1998;18(3):841–853. 10.1523/jneurosci.18-03-00841.1998.

Sultan ZW, Najac M, Raman IM. Control of action potential afterdepolarizations in the inferior olive by inactivating A-type currents through KV4 channels. The Journal of Physiology. 2024;10.1113/jp286818.

Suter KJ, Jaeger D. Reliable control of spike rate and spike timing by rapid input transients in cerebellar stellate cells. Neuroscience. 2004;124(2):305–317. 10.1016/j.neuroscience.2003.11.015.

Szapiro G, Barbour B. Multiple climbing fibers signal to molecular layer interneurons exclusively via glutamate spillover. Nature Neuroscience. 2007;10(6):735–742. 10.1038/nn1907.

Teruyama R, Armstrong WE. Changes in the Active Membrane Properties of Rat Supraoptic Neurones During Pregnancy and Lactation. Journal of Neuroendocrinology. 2002;14(12):933–944. 10.1046/j.1365-2826.2002.00844.x.

Tran-Van-Minh A, Abrahamsson T, Cathala L, DiGregorio DA. Differential Dendritic Integration of Synaptic Potentials and Calcium in Cerebellar Interneurons. Neuron. 2016;91(4):837–850. 10.1016/j.neuron.2016.07.029.

Traub RD, Wong RK, Miles R, Michelson H. A model of a CA3 hippocampal pyramidal neuron incorporating voltage-clamp data on intrinsic conductances. Journal of Neurophysiology. 1991;66(2):635–650. 10.1152/jn.1991.66.2.635.

Wang XJ. Probabilistic Decision Making by Slow Reverberation in Cortical Circuits. Neuron. 2002;36(5):955–968. 10.1016/s0896-6273(02)01092-9.

Watanabe J, Beck C, Kuner T, Premkumar LS, Wollmuth LP. DRPEER: A Motif in the Extracellular Vestibule Conferring High Ca2+Flux Rates in NMDA Receptor Channels. The Journal of Neuroscience. 2002;22(23):10209–10216. 10.1523/jneurosci.22-23-10209.2002.

White G, Lovinger DM, Weight FF. Transient low-threshold Ca2+ current triggers burst firing through an afterdepolarizing potential in an adult mammalian neuron. Proceedings of the National Academy of Sciences. 1989;86(17):6802–6806. 10.1073/pnas.86.17.6802.

Williams SR, Stuarty GJ. Mechanisms and consequences of action potential burst firing in rat neocortical pyramidal neurons. The Journal of Physiology. 1999;521(2):467–482. 10.1111/j.1469-7793.1999.00467.x.

Wong RK, Prince DA. Afterpotential generation in hippocampal pyramidal cells. Journal of Neurophysiology. 1981;45(1):86–97. 10.1152/jn.1981.45.1.86.

Wu WW, Chan CS, Disterhoft JF. Slow After-hyperpolarization Governs the Development of NMDA Receptor–Dependent Afterdepolarization in CA1 Pyramidal Neurons During Synaptic Stimulation. Journal of Neurophysiology. 2004;92(4):2346–2356. 10.1152/jn.00977.2003.

Xu J, Clancy CE. Ionic Mechanisms of Endogenous Bursting in CA3 Hippocampal Pyramidal Neurons: A Model Study. PLoS ONE. 2008;3(4):e2056. 10.1371/journal.pone.0002056.

Yi G, Cui J, Bai R. Effects of dendritic properties on the correlations in ionic channels emerging from firing rate homeostasis: a two-compartment modeling study. Cognitive Neuro-dynamics. 2025;19(1). 10.1007/s11571-025-10297-z.

Yoshimura M, Polosa C, Nishi S. Afterdepolarization mechanism in the in vitro, cesium-loaded, sympathetic preganglionic neuron of the cat. Journal of Neurophysiology. 1987;57(5):1325–1337. 10.1152/jn.1987.57.5.1325.

Yue C, Yaari Y. KCNQ/M Channels Control Spike Afterdepolarization and Burst Generation in Hippocampal Neurons. The Journal of Neuroscience. 2004;24(19):4614–4624. 10.1523/jneurosci.0765-04.2004.

Yue C, Yaari Y. Axo-Somatic and Apical Dendritic Kv7/M Channels Differentially Regulate the Intrinsic Excitability of Adult Rat CA1 Pyramidal Cells. Journal of Neurophysiology. 2006;95(6):3480–3495. 10.1152/jn.01333.2005.

