## Supplemental Material for "Dendritic Interaction of Timescales in Afterdepolarization Potentials and Nonmonotonic Spike-adding"

The ball-and-stick CSC model consists of a somatic compartment and a dendritic cable, discretized into 1 to $Nseg$ compartments. It is given by

$$\begin{aligned}
C\dot{V}_s &= -I_{Na,s} - I_{K,s} - I_{Leak,s} - I_{A,s} - I_{T,s} \\
&- I_{K_{Ca,s}} - I_{HVA,s} + I_{d1,s} \\
C\dot{V}_1 &= -I_{Na,1} - I_{K,1} - I_{Leak,1} - I_{A,1} - I_{T,1} \\
&- I_{K_{Ca,1}} - I_{HVA,1} + I_{s,1} + I_{1,2} \\
C\dot{V}_i &= -I_{Na,i} - I_{K,i} - I_{Leak,i} - I_{A,i} - I_{T,i} \\
&- I_{K_{Ca,i}} - I_{HVA,i} + I_{i-1,i} + I_{i,i+1} \\
&\text{for } i \in [2, Nseg] \\
C\dot{V}_{Nseg} &= -I_{Na,Nseg} - I_{K,i} - I_{Leak,Nseg} \\
&- I_{A,Nseg} - I_{T,Nseg} - I_{K_{Ca,Nseg}} \\
&- I_{HVA,Nseg} + I_{Nseg-1,Nseg}
\end{aligned} \tag{8}$$

The resistive coupling from cylindrical compartments $\mu'$ to $\mu$ due to Ohm's Law is

$$g_{\mu,\mu'} = \frac{a_\mu a_{\mu'}}{a_L L_\mu (L_\mu a_{\mu'}^2 + L_{\mu'} a_\mu^2)},$$

where $a$ is the radius and $L$ is the length of the respective compartments Dayan et al. (2001).

Thus, for compartments of the same dimensions, such as those in the discretized cable of uniform diameter, the conductance and current from adjacent compartment $\mu$ to $\mu'$ is

$$\begin{aligned}
g_{\mu,\mu'} &= \frac{r_L L}{\pi a^2} \\
I_{\mu,\mu'} &= g_{\mu,\mu'}(V_{\mu'} - V_\mu)
\end{aligned}$$

**Table S1** Parameter values of ionic currents for the ball-and-stick CSC model. Conductances of the somatic and dendritic compartments of the two-compartment are shown separately.

| | $g_{soma}$ $\mu S/cm^2$ | $g_{dendrites}$ $\mu S/cm^2$ |
|---|---|---|
| $I_{Na}$ | 3.4 | — |
| $I_K$ | 10.0112 | 10.2388 |
| $I_A$ | 6.8491 | 5.3509 |
| $I_T$ | 0.2834 | 0.1670 |
| $I_{HVA}$ | 0.02047 | 0.05953 |
| $I_L$ | 0.07407 | 0.07407 |
| $I_{K(Ca)}$ | 0.5505 | 0.4495 |

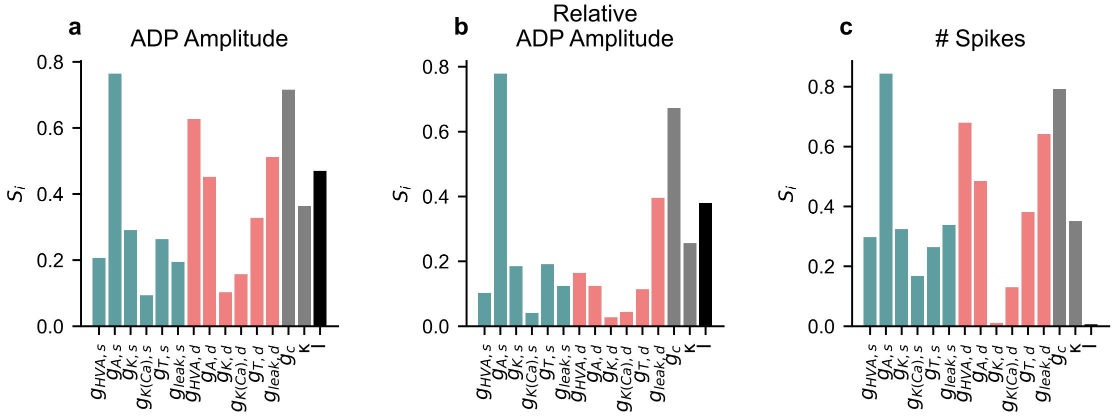


**Fig. S1** Sensitivity analysis with somatic input. Variance-based sensitivity analysis resulting the Sobol indices relating to ADP amplitude (a), relative ADP amplitude (b) and the number of evoked APs (c) during somatic step current input for somatic and dendritic model conductances, as well as for coupling conductance ($g_c$), proportion of somatic to total surface area ($\kappa$) and step current magnitude (I)

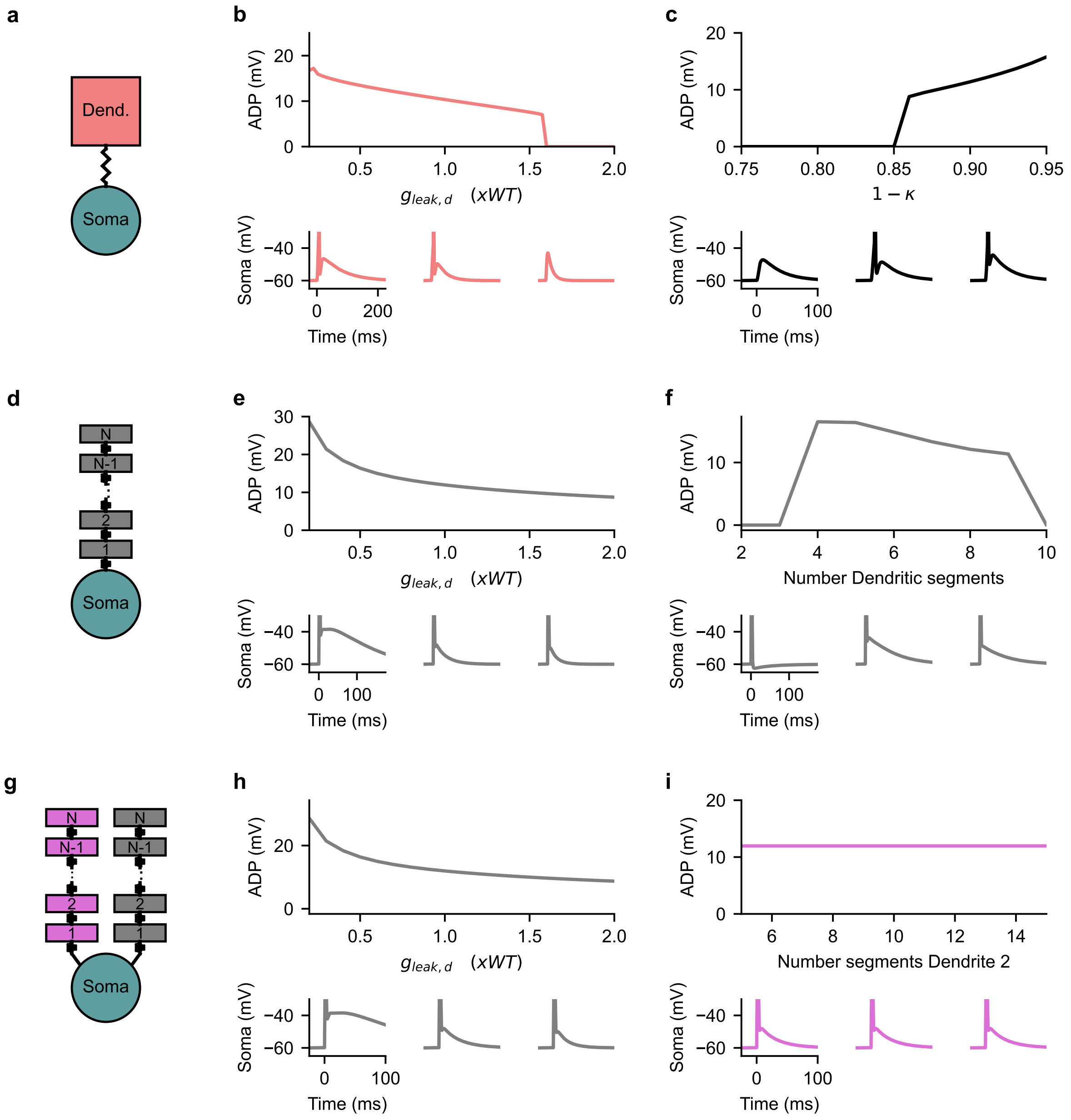


**Fig. S2** Effect of Morphology on afterdepolarizations. (a) Schematic of the two compartmental model receiving a dendritic step current input. This two compartment model was used to study (b) the effect of scaling the dendritic leak conductance ($g_{leak,d}$) from its default value (WT) on the amplitude of the ADP, as well as analyze (c) the effect of increasing the dendritic surface area, i.e, increasing $1-\kappa$, on the amplitude of ADPs. (d-f) The equivalent effects of dendritic leak conductance and dendritic length on ADP amplitude using the ball-and-stick model. (g-i) The equivalent effects of dendritic leak conductance and dendritic length on ADP amplitude using the ball-and-stick model but with a second cable representing another dendritic branch.

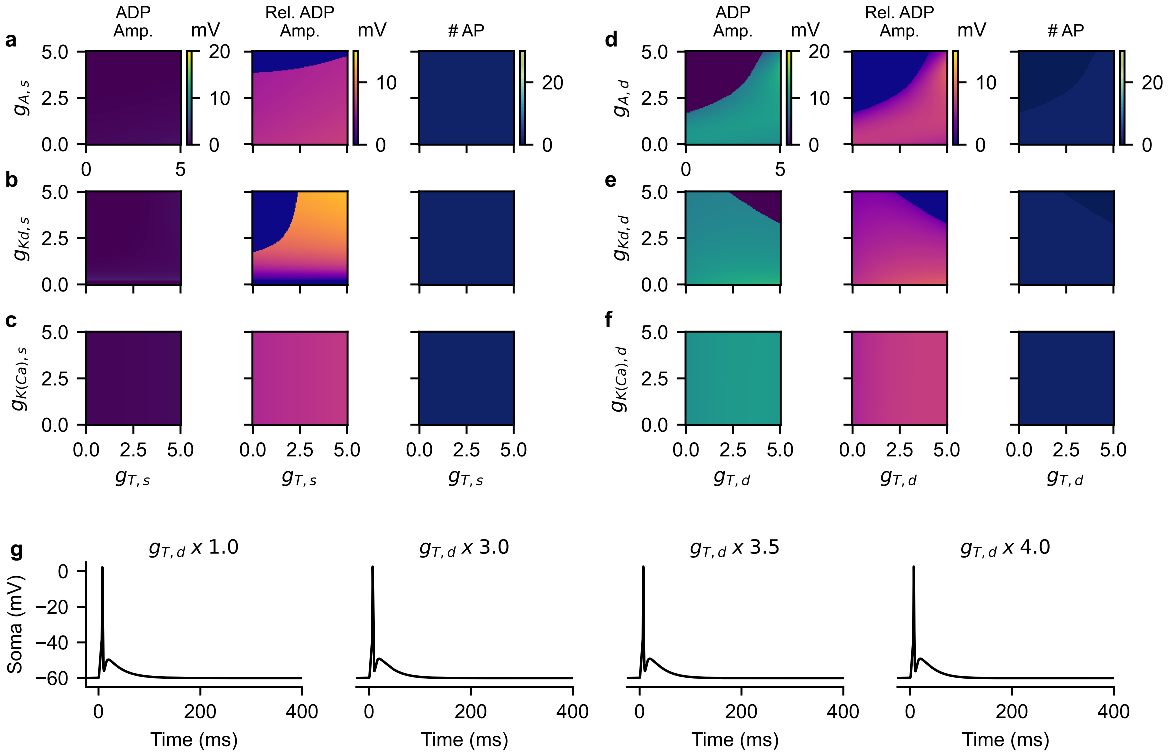


**Fig. S3** Dendritic T-type $Ca^{2+}$ and $K^+$ currents do not evoke spike-adding with step current input. The effect of altering somatic A-type $K^+$ (a; $g_{A,s}$), Kdr (b; $g_{Kd,s}$) and K(Ca) (c; $g_{K(Ca),s}$) current conductances in combination with somatic T-type $Ca^{2+}$ ($g_{T,s}$) current conductance during 3 ms step current input (20 $\mu A/cm^2$) on ADP amplitude, relative ADP amplitude and the number of spikes evoked. The effect of altering dendritic A-type $K^+$ (d; $g_{A,d}$), Kdr (e; $g_{Kd,d}$) and K(Ca) (f; $g_{K(Ca),d}$) and dendritic T-type $Ca^{2+}$ ($g_{T,d}$) during step current input on ADP amplitude, relative ADP amplitude and the number of spikes evoked. The somatic voltage responses (g) demonstrate the minimal effect of increasing dendritic HVA $Ca^{2+}$ conductance in d-f. All conductance values are reported in fold change from its default value.

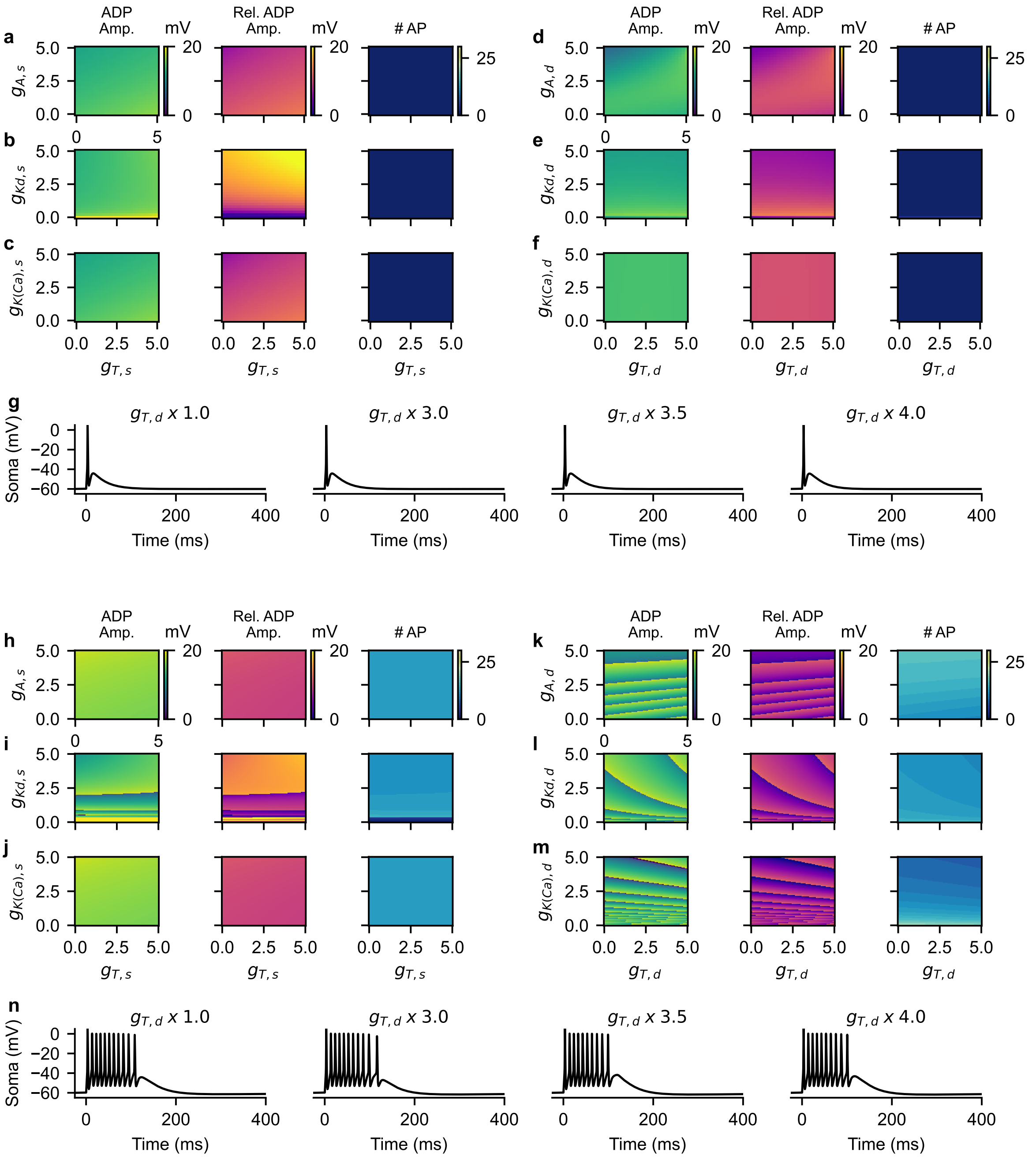


**Fig. S4** Dendritic T-type $Ca^{2+}$ and $K^+$ currents do not modulate spike-adding with AMPA current inputs but do with NMDA current inputs. The effect of altering somatic A-type $K^+$ (a; $g_{A,s}$), Kdr (b; $g_{Kd,s}$) and K(Ca) (c; $g_{K(Ca),s}$) current conductances in combination with somatic T-type $Ca^{2+}$ ($g_{T,s}$) current conductance on ADP amplitude, relative ADP amplitude and the number of evoked APs in the presence of AMPA current input ($g_{AMPA} = 10\ \mu S/cm^2$). The effect of altering dendritic A-type $K^+$ (d; $g_{A,d}$), Kdr (e; $g_{Kd,d}$) and K(Ca) (f; $g_{K(Ca),d}$) in concert with dendritic T-type $Ca^{2+}$ ($g_{T,d}$) on ADP amplitude, relative ADP amplitude and the number of evoked APs during the same AMPA current input. The somatic voltage responses (G) demonstrate the lack of spike-adding with increasing dendritic T-type $Ca^{2+}$ conductance in d-f with AMPA current input. After the addition of an NMDA current to the AMPA current input ($g_{NMDA} = 2\ \mu S/cm^2$), the effects of somatic A-type $K^+$ (h; $g_{A,s}$), Kdr (i; $g_{Kd,s}$) and K(Ca) (j; $g_{K(Ca),s}$) in relation to somatic T-type $Ca^{2+}$ ($g_{T,s}$) as well as dendritic A-type $K^+$ (k; $g_{A,d}$), Kdr (l; $g_{Kd,d}$) and K(Ca) (m; $g_{K(Ca),d}$) during altered dendritic T-type $Ca^{2+}$ ($g_{T,d}$) were assessed. The somatic voltage responses (N) to an AMPA and NMDA current input evokes numerous action potentials across dendritic T-type $Ca^{2+}$ conductance in d-f. All conductance values are reported in fold change from its default value.

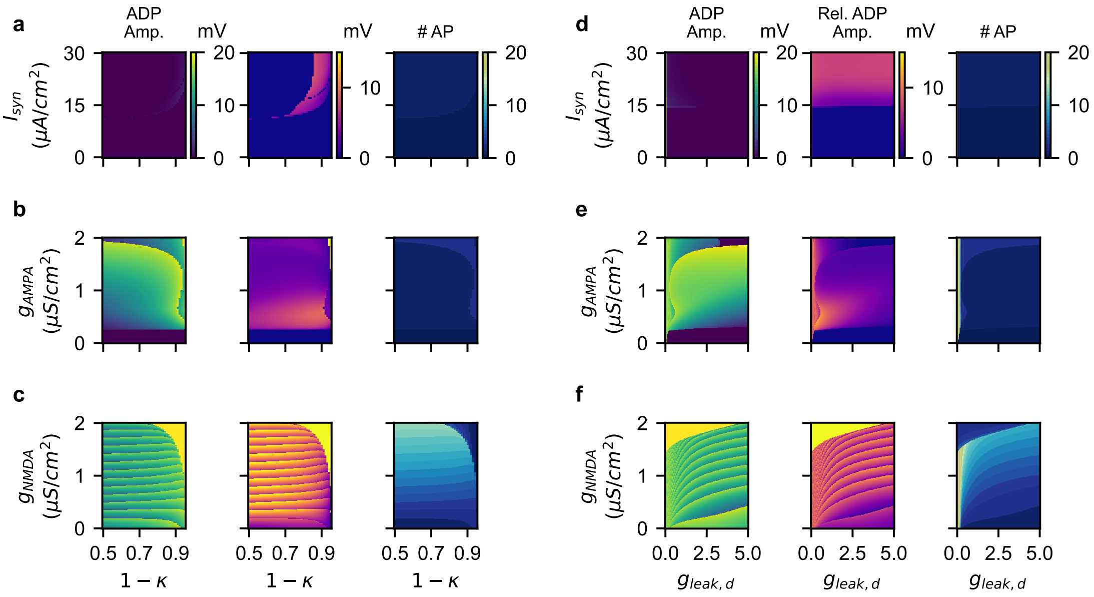


**Fig. S5** Effects of coupling and dendritic leak current with somatic input. The effect of dendritic length, implemented in the two compartmental model as the ratio of dendritic to total surface area $(1 - \kappa)$, as well as step current input magnitude ($I_{syn}$; a), AMPA current conductance ($g_{AMPA}$; b) and NMDA current conductance ($g_{NMDA}$; c) on ADP amplitude, relative ADP amplitude and number of evoked APs. The effect of dendritic leak current magnitude is modulated by altering the dendritic leak conductance $g_{leak,d}$ (in fold change from its default value) and evaluated in combination with step current input magnitude (d), AMPA current conductance (e) and NMDA current conductance (f). NMDA synaptic input occurred in the presence of an AMPA current with fixed conductance (10 $\mu$S/cm$^2$
